# Widespread fungal-bacterial competition for magnesium enhances antibiotic resistance

**DOI:** 10.1101/2023.10.25.563990

**Authors:** Yu-Ying Phoebe Hsieh, Wanting Wendy Sun, Janet M. Young, Robin Cheung, Deborah A. Hogan, Ajai A. Dandekar, Harmit S. Malik

**Author notes:** Correspondence should be addressed to: Yu-Ying Phoebe Hsieh, 1100 Fairview Avenue N. A2-205, Seattle WA 98109. equal contribution (Ajai A. Dandekar, Harmit S. Malik).

## Abstract

Fungi and bacteria coexist in many polymicrobial communities, yet the molecular basis of their interactions remains poorly understood. Using unbiased genomic approaches, we discover that the fungus *Candida albicans* sequesters essential Mg^2+^ ions from the bacterium *Pseudomonas aeruginosa*. In turn, the bacterium competes using a Mg^2+^ transporter, MgtA. We show that Mg^2+^ sequestration by fungi is a general mechanism of antagonism against gram-negative bacteria. But the resultant Mg^2+^ limitation enhances bacterial resistance to polymyxin antibiotics like colistin, which target gram-negative bacterial membranes. Experimental evolution reveals that bacteria in co-culture with fungi become phenotypically, but not genetically, resistant to colistin; antifungal treatment renders resistant bacteria from co-cultures to become colistin-sensitive. Fungal-bacterial nutritional competition may thus profoundly impact treatments of polymicrobial infections with antibiotics of last resort.

**One Sentence Summary:** Magnesium sequestration by fungi lowers bacterial fitness but enhances antibiotic resistance.

## Main Text

Fungal-bacterial interactions are pivotal in shaping the success and composition of polymicrobial communities in various environments (*1, 2*). Cooperation and competition between fungi and bacteria in host-associated communities, including animal gastrointestinal tracts and chronically infected tissues (*3*), can significantly impact both microbial and host fitness. Polymicrobial infections are more recalcitrant to treatment by either antibiotic or antifungal agents (*4*). The fungus *Candida albicans* and the bacterium *Pseudomonas aeruginosa* are opportunistic human pathogens that infect chronic wounds (*5*) and the airways of people with cystic fibrosis (*6, 7*). Previous studies have identified strategies that *C. albicans* and *P. aeruginosa* use to antagonize each other, including bacterial toxins that target fungal hyphae (*8, 9*), interference of quorum sensing regulations (*10, 11*), and competition for limited resources like iron (*12*). Most studies have focused on anti-fungal strategies imposed by bacteria. However, fungal strategies for competing with bacteria and whether these mechanisms are species-specific remain unknown. Another central question is how fungal-bacterial competition influences the evolution of drug resistance. This knowledge could be leveraged to predict resistance development in polymicrobial infections and improve therapies to cure infectious diseases. We aimed to identify strategies fungi use to antagonize bacteria, potential bacterial counterstrategies, and the consequences of such fungal-bacterial competition on antibiotic resistance.

## Results

To identify strategies fungi use to antagonize bacteria, we first demonstrated that *C. albicans* impairs the fitness of *P. aeruginosa* (strain PAO1) in brain heart infusion (BHI) broth (**Fig. S1A**), a widely used medium for studying human microbial pathogens. To understand the physiological basis for this impairment, we performed RNA-seq analyses on *P. aeruginosa* following eight hours of co-culture with *C. albicans* (strain SC5314) relative to bacteria-only conditions (monoculture); bacterial fitness at this time point was nearly identical between the two conditions (**Fig. S2**). Our analyses revealed that 145 *P. aeruginosa* genes were up-regulated by at least four-fold in co-culture relative to monoculture conditions, including those related to TonB-dependent substrate transport, siderophore synthesis, and RNA polymerase sigma factor 70 (**Fig. S3A and Table S1**). We also found that 134 genes, including those for Type VI secretion system, co-factor biosynthesis, and energy generation, were down-regulated by at least four-fold in co-culture (**Fig. S3B and Table S1**). These changes in gene expression suggest that *P. aeruginosa* prioritized nutrient uptake while minimizing energy production in co-culture.

We hypothesized that *P. aeruginosa* might rely on defense genes to protect itself during co-culture with *C. albicans*; loss of such defense genes would further impair *P. aeruginosa* fitness, specifically under co-culture conditions (*13*). To identify genes important for bacterial fitness in the presence of fungi, we used a transposon-insertion sequencing (Tn-seq) approach (*14–16*) to conduct a genome-wide fitness screen in bacteria. We cultured a pool of 10^5^ unique *P. aeruginosa* transposon-insertion (Tn) mutants (*14*) for ten generations, either in monoculture or in co-culture with *C. albicans.* We then quantified the fitness value of each Tn mutant by comparing the number of reads of each transposon mutant in both conditions (**Fig. 1A**). Our findings revealed that Tn insertions in eight *P. aeruginosa* genes significantly reduced bacterial fitness in co-culture relative to monoculture conditions (**Fig. 1B and C**). Three adjacent genes – *PA4824*, *PA4825*, and *PA4826* – showed the most significant fitness loss in co-culture (**Fig. 1B**). These three genes were also the only overlap between our Tn-seq and RNA-seq analyses, with *PA4824* and *PA4825* showing a 32-fold increase in expression during co-culture conditions (**Fig. S3C**). Of these three genes, only *PA4825* has been functionally characterized; it encodes a magnesium transporter known as MgtA. In addition to the eight genes in which Tn insertions decreased fitness (defense genes), our Tn-seq analyses identified 18 genes whose loss enhanced bacterial fitness in co-culture (**Table S2**), suggesting these genes either become ‘dispensable’ or incur a fitness cost in co-culture conditions.

**Figure 1.**
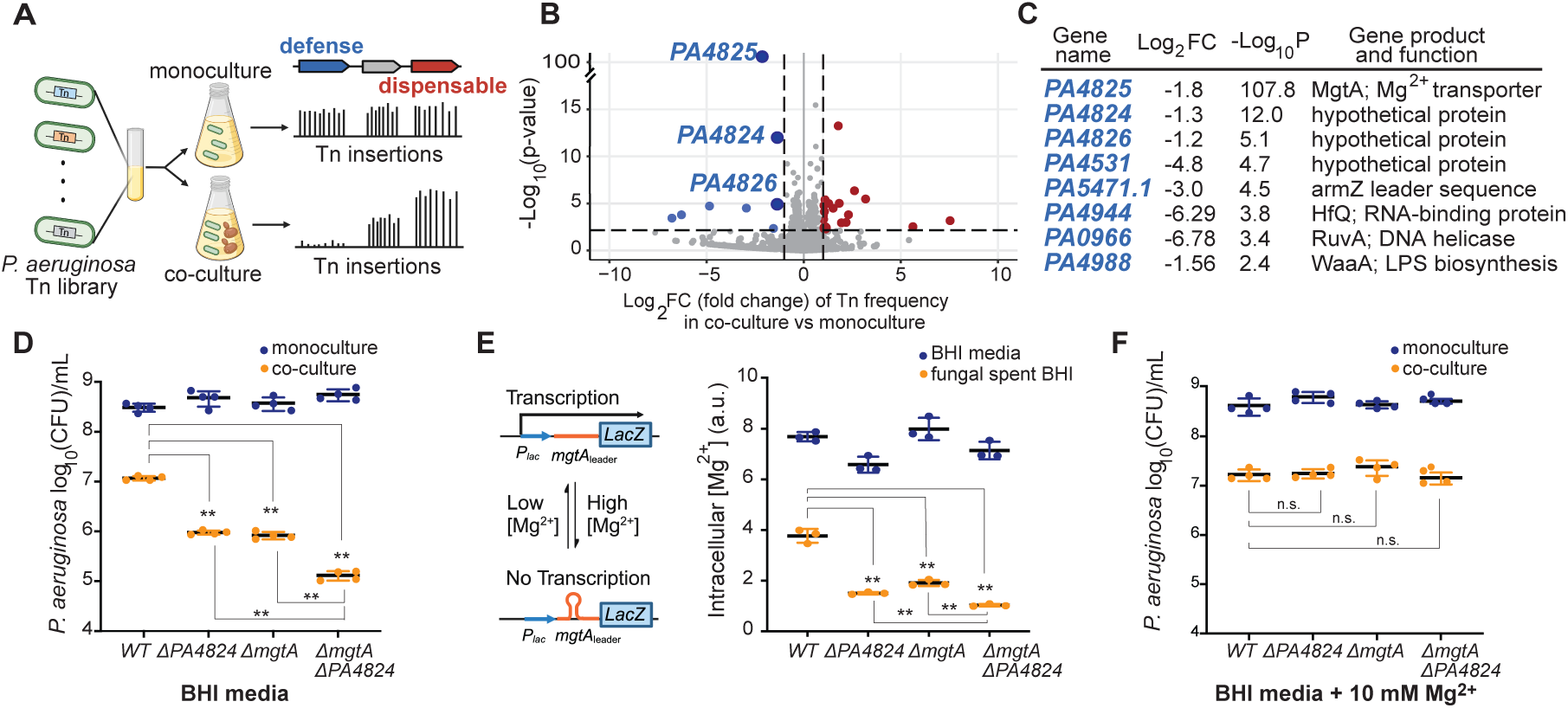
*P. aeruginosa* is suppressed by *C. albicans*-mediated Mg^2+^ sequestration. **(A)** A pool of *P. aeruginosa* Tn mutants was grown in BHI media only or in co-culture with *C. albicans* in BHI. Quantification of Tn mutant frequencies (see Methods) revealed genes whose loss impaired *P. aeruginosa* fitness (blue; defense genes) and genes whose loss conferred a fitness advantage in co-culture with *C. albicans* (red; dispensable genes). **(B)** Tn-seq volcano plot (see Methods) shows *P. aeruginosa* genes that were important for defense or dispensable in co-culture with *C. albicans*. X-axis indicates fold change, while Y-axis indicates *p*-value after correcting for multiple testing. We used log_2_ fold change > 2 and adjusted *p*-value < 0.1 as statistical cut-offs. **(C)** Candidate defense gene identified by Tn-seq. **(D)** Fitness of *P. aeruginosa* single deletion Δ*PA4824* or Δ*mgtA* mutants was impaired relative to WT *P. aeruginosa* in co-culture with *C. albicans* (orange) but not monoculture in BHI media (blue). The fitness of double deletion Δ*PA4824* Δ*mgtA* mutant was even further impaired than single deletion mutants. Colony-forming units (CFUs) of *P. aeruginosa* were enumerated by serially diluting cultures on LB+Nystatin. **(E)** (left panel) Intracellular Mg^2+^ in *P. aeruginosa* is measured using an RNA sensor. When the intracellular Mg^2+^ level is sufficient, *mgtA* 5’UTR forms a secondary structure that blocks downstream transcription. In limiting Mg^2+^, this structure is resolved, and transcription of a β-galactosidase reporter is restored. (right panel) Intracellular Mg^2+^ levels, measured in β-galactosidase units, of single deletion mutants lacking *PA4824* or *mgtA* were lower than the WT stain and even lower in a double deletion Δ*PA4824* Δ*mgtA* mutant in *C. albicans*-spent BHI (orange), but not BHI media alone (blue). (**F**) Co-culture-specific fitness loss of the *P. aeruginosa* single or double deletion mutants relative to WT strains was restored by Mg^2+^ supplementation (10mM) in BHI in co-culture (orange). Mean ± std of three biological replicates is shown in panels D-F. (** *p* < 0.01 unpaired two-tailed Student’s *t*-test used; n.s. indicates no significance).

### Magnesium is an axis of nutritional competition between *P. aeruginosa* and *C. albicans*

We chose to focus on the contributions of the *PA4824*, *PA4825 (mgtA)*, and *PA4826* genes during co-culture. We engineered gene deletion mutants and measured the fitness of these mutants relative to a fluorescently labeled wild-type (WT) strain in either monoculture or co-culture conditions using fitness competition assays, which resemble the pooled Tn-seq bulk selection conditions (**Fig. S4A**). Our experiments showed that the relative fitness of the Δ*PA4824* or Δ*PA4825* (Δ*mgtA)* mutants, but not Δ*PA4826,* was ∼10% lower than the WT in co-culture with *C. albicans* (**Fig. S4B**). These mutants had no fitness defect under monoculture conditions (**Fig. S4B**). Co-culture colony formation assays also confirmed our findings. Deletion of either Δ*PA4824* or Δ*mgtA* led to a 10-fold reduction in colony forming units (CFU) of *P. aeruginosa* in co-culture relative to monoculture (**Fig. 1D**). Moreover, a double deletion Δ*PA4824* Δ*mgtA* mutant was even more significantly impaired than single deletion mutants (**Fig. 1D**). These results suggest that the protein products encoded by *PA4824* and *mgtA* are independently required for optimal *P. aeruginosa* fitness under co-culture conditions, but *PA4826* is not; its identification in our Tn-seq analyses may have been due to insertion mutant effects on adjacent genes.

Our finding that *P. aeruginosa PA4825 (mgtA)* is required for full bacterial viability in co-culture led us to hypothesize that *C. albicans* depletes the vital cation Mg^2+^ in co-culture. Mg^2+^ depletion would make *mgtA* essential and lower intracellular Mg^2+^ levels in the Δ*mgtA* mutant. To test this hypothesis, we employed a Mg^2+^ genetic sensor (*17*) from *Salmonella enterica* serovar Typhimurium. *S.* Typhimurium *mgtA* expression levels are controlled by its 5’ untranslated region (5’UTR), which acts as a ribo-switch, adopting different stem-loop structures depending on the Mg^2+^ levels and regulating the transcription of *mgtA* (*17*). We showed that this *S.* Typhimurium Mg^2+^ sensor can accurately detect changes in intracellular *P. aeruginosa* Mg^2+^ levels (**Fig. S5**). Using this sensor, we found that the intracellular Mg^2+^ levels in the WT *P. aeruginosa* are reduced by two-fold in fungal spent media (*i.e.,* derived from the filtrate of *C. albicans* cultures to simulate Mg^2+^ depletion) compared to in fresh BHI media (**Fig. 1E**). Loss of *mgtA* further reduced bacterial intracellular Mg^2+^ levels in the fungal spent media, but not in monoculture conditions (**Fig. 1E**). Surprisingly, loss of *PA4824* also reduced intracellular Mg^2+^ levels in fungal spent media compared to the WT strain, with a double deletion Δ*PA4824* Δ*mgtA* mutant further reducing intracellular Mg^2+^ levels compared to single gene deletion mutants (**Fig. 1E**). These experiments support the hypothesis of fungal-imposed Mg^2+^ sequestration and demonstrate that *mgtA* and *PA4824* are independently required to restore intracellular Mg^2+^ levels and *P. aeruginosa* fitness in co-culture with *C. albicans*.

Our findings suggest that *PA4824* is important for Mg^2+^ uptake from the environment. Although *PA4824* remains functionally uncharacterized, structural predictions based on Alphafold (*18*) revealed that it encodes a transmembrane protein with a distinctive β-barrel structure commonly associated with proteins involved in ion transport or nutrient uptake (**Fig. S6A**). The core of PA4824 is composed of hydrophilic and negative-charged residues (**Fig. S6B and C**), suggesting it might transport positive-charged molecules, potentially acting as a novel Mg^2+^ transporter. In addition to MgtA (and potentially PA4824), *P. aeruginosa* uses two other Mg^2+^ transporters, CorA and MgtE, to obtain Mg^2+^ from environments (*19*). However, neither of these transporters was identified in our Tn-seq or RNA-seq analyses (**Fig. S7A**). Indeed, we found that *corA* or *mgtE* loss-of-function mutants did not alter *P. aeruginosa* fitness in co-culture conditions (**Fig. S7B**). This indicates that MgtA is a bacterial Mg^2+^ transporter that is highly induced under low Mg^2+^ conditions (*20*) and required to overcome fungal sequestration of Mg^2+^.

If *C. albicans* consumes or sequesters Mg^2+^ from *P. aeruginosa*, we hypothesized that supplementing cultures with Mg^2+^ might rescue the viability reduction of Δ*mgtA* and Δ*PA4824* mutants relative to the WT strain. Indeed, supplemental Mg^2+^ (10 mM) was sufficient to restore the fitness of Δ*mgtA,* Δ*PA4824,* and Δ*PA4824* Δ*mgtA* double deletion mutants to the WT level in co-culture conditions (**Fig. 1F**). To investigate whether Mg^2+^ could explain fitness effects of other candidate genes revealed by the Tn-seq analyses, we repeated our Tn-seq experiments in monoculture versus co-culture conditions but with supplemental Mg^2+^ (10 mM). To our surprise, we found that Mg^2+^ supplementation ameliorated the fitness effects of most of the candidate genes identified in our initial Tn-seq analyses (**Fig. S8A**), including all eight defense genes and 17 of 19 dispensable genes crucial to fundamental cellular processes, including cell division, pyrimidine synthesis, and energy production. As enzymes in these essential processes rely on Mg^2+^, we hypothesized that they might become too costly to maintain in low Mg^2+^ co-culture conditions. Only two genes (*PA4029* and *PA5484)* remained marginally essential in co-culture even under supplemented Mg^2+^ conditions (**Fig. S8A**). Notably, Mg^2+^ supplementation only affected the expression of *PA4824*-*PA4826*, but not other candidate genes (**Fig. S8B**), suggesting that Mg^2+^ levels affect their functions post-transcriptionally. Thus, our finding underscores the influential role of Mg^2+^ competition in mediating the impact of most candidate genes on overall bacterial fitness in co-culture conditions. We note, however, that Mg^2+^ supplementation did not fully restore the fitness of *P. aeruginosa* WT cells in co-culture to monoculture fitness levels (**Fig. 1F**), indicating that additional, Mg^2+^-independent, axes of antagonism exist (*21–23*).

Nutritional competition for important metal ions, such as iron, zinc, and manganese, is a common axis of competition among microbes and between microbes and vertebrate hosts (*24, 25*). However, competition for the vital Mg^2+^ cation is a novel finding. We reasoned that our finding, despite previous efforts to identify determinants of competition between *C. albicans* and *P. aeruginosa*, might reflect different Mg^2+^ levels in the various media conditions used for co-culture. Indeed, we found that MgtA is required for *P. aeruginosa* fitness in co-culture with *C. albicans* in BHI and Tryptic Soy Broth (TSB) media, but not in Yeast Extract-Peptone-Dextrose (YPD) or synthetic cystic fibrosis medium (SCFM) (*26*) (**Fig. S9A**). Mg^2+^ levels in BHI and TSB were lower (100-140μM) than in YPD or SCFM (240-390μM) (**Fig. S9B**). As a result, fungal filtrates derived from BHI and TSB exhibited the lowest levels of Mg^2+^ (**Fig. S9B**), leading to a significant reduction of intracellular Mg^2+^ in *P. aeruginosa* (**Fig. S9C**). These results indicated that BHI represents a low Mg^2+^ condition, leading to our discovery of Mg^2+^ competition. Consistent with these data, supplementation of BHI with greater than 300μM Mg^2+^ was sufficient to relieve the fitness defect of the *P. aeruginosa* Δ*mgtA* mutant (**Fig. S9D**). Thus, fungal-mediated Mg^2+^ competition against bacteria is most likely to occur in conditions where local Mg^2+^ is scarce, such as in biofilms (*27*), polymicrobial communities where microbes overproduce cation-chelating agents such as citrate (*28*), infected tissues where host immune cells rigorously control Mg^2+^ (*29*), or in magnesium-deficient hosts (*30, 31*).

### Widespread Mg^2+^ competition between fungi and gram-negative bacteria

We next assessed whether Mg^2+^ competition is a common means of fungal antagonism against bacteria. *Escherichia coli* and *S.* Typhimurium are gram-negative bacteria that co-exist with *C. albicans* in the human gut (*32*). *E. coli* encodes a single *mgtA* ortholog (like *P. aeruginosa*) whereas *S.* Typhimurium encodes two paralogs: *mgtA* and *mgtB*. We found that either a single deletion Δ*mgtA* in *E. coli* or a double deletion Δ*mgtA* Δ*mgtB* in *S.* Typhimurium severely reduced bacterial fitness in co-culture with *C. albicans*, but bacterial fitness could be restored via Mg^2+^ supplementation (**Fig. 2A and B**). Next, we expanded our analyses to three additional fungal species that co-exist with *P. aeruginosa* in polymicrobial infections (*33*): *Candida tropicalis*, *Candida parapsilosis*, *Candida glabrata*, as well as a non-pathogenic species, *Saccharomyces cerevisiae*. In each case, we found that the *P. aeruginosa* Δ*mgtA* mutant had reduced fitness relative to WT cells in fungal co-culture (**Fig. 2C**), which could be restored by Mg^2+^ supplementation (**Fig. 2D**). Our findings confirm that competition for Mg^2+^ is a widespread mode of fungal antagonism against gram-negative bacteria and that MgtA, a gram-negative bacteria-specific Mg^2+^ transporter (*34*), plays an important role in defense against fungal-mediated Mg^2+^ competition.

**Figure 2.**
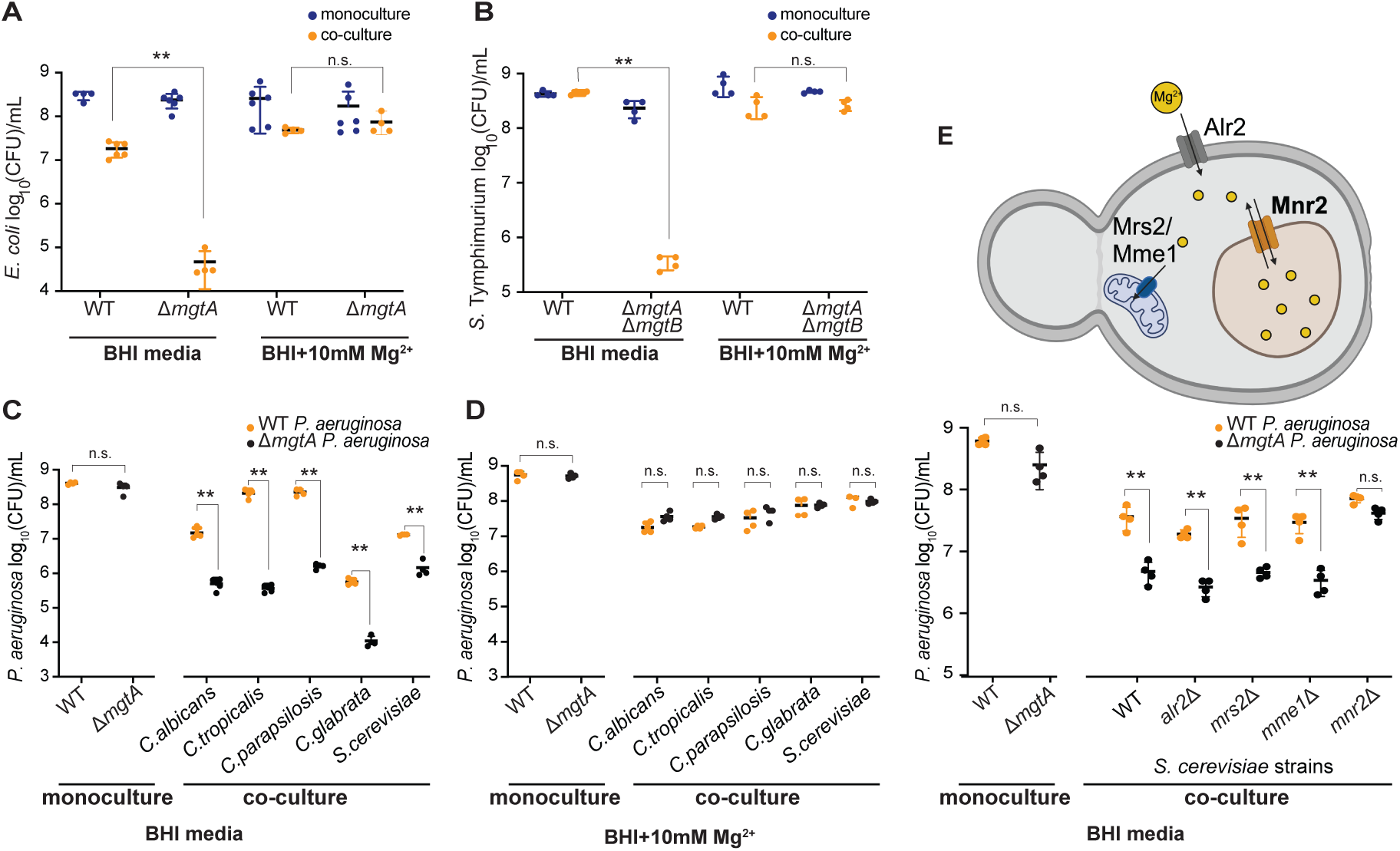
Mg^2+^ competition is widespread between multiple fungal and gram-negative bacterial species. **(A)** Fitness of the *E. coli* Δ*mgtA* mutant was impaired relative to the WT strain in co-culture with *C. albicans* (orange) but not in monoculture (blue) in BHI media. The fitness of the Δ*mgtA* strain was restored in BHI media supplemented with 10 mM Mg^2+^. **(B)** Fitness of the double deletion Δ*mgtA* Δ*mgtB* mutant *S. Typhimurium* strain was impaired relative to WT in co-culture with *C. albicans* (orange) but not in monoculture in BHI media (blue). **(C)** Fitness of the Δ*mgtA* mutant *P. aeruginosa* strain (black) was impaired relative to WT (orange) in co-culture with various fungal species (*C. albicans, C. tropicalis, C. parapsilosis, C. glabrata, S. cerevisiae*) in BHI. **(D)** The functions and locations of proposed Mg^2+^ transporters in *S. cerevisiae* are illustrated (top panel); Alr2 localizes to the plasma membrane, Mrs2 and Mme1 to the mitochondria, and Mnr2 to the vacuole. The fitness of the *P. aeruginosa* Δ*mgtA* mutant (black) was similar to the WT strain (orange) in monoculture. However, the relative fitness of *P. aeruginosa* Δ*mgtA* was impaired in co-culture with either of three *S. cerevisiae* mutant strains – Δ*alr2,* Δ*mrs2, or* Δ*mme1* – just as it is in co-culture with WT *S. cerevisiae*. In contrast, the relative fitness of *P. aeruginosa* Δ*mgtA* was restored in co-culture with *S. cerevisiae* Δ*mnr2* strain. Mean ± std of three biological replicates is shown in panels A-E. (** *p* < 0.01 unpaired two-tailed Student’s *t*-test used; n.s. indicates no significance).

How do fungi impose competition for Mg^2+^? We investigated whether loss of *S. cerevisiae* genes involved in Mg^2+^ uptake or sequestration can restore the fitness of bacterial Δ*mgtA* mutants. *S. cerevisiae* cells store 80% of cellular Mg^2+^ in their vacuole (*35*). We found that the fitness of *P. aeruginosa* Δ*mgtA* mutant was restored to WT levels in co-culture with *S. cerevisiae* lacking Mnr2 (*36*), previously proposed to be a vacuolar Mg^2+^ transporter (**Fig. 2E**). In contrast, *P. aeruginosa* Δ*mgtA* fitness was not restored by competition with *S. cerevisiae* mutants lacking Alr2, a cell-wall-associated Mg^2+^ transporter used to acquire environmental Mg^2+^ into the cytosol (*37*), or those lacking Mrs2/Mme1, mitochondria-specific Mg^2+^ transporters (**Fig. 2E**). Thus, we hypothesize that fungal vacuolar Mg^2+^ sequestration might broadly mediate Mg^2+^ antagonism against gram-negative bacteria.

### Fungal Mg^2+^ sequestration protects bacteria from polymyxin antibiotics

Mg^2+^ is required for many fundamental cellular processes, acting as a key player in neutralizing negatively charged biomolecules and serving as a cofactor for many enzymes (*19*). Mg^2+^ is also important for stabilizing the outer membrane structure of gram-negative bacteria by binding to negatively charged lipopolysaccharides (LPS) present on the bacterial cell membrane (*38*). These Mg^2+^-stabilized LPS moieties can be targeted by polymyxins (*39*), last-resort antibiotics for treating multidrug-resistant bacterial infections (*40, 41*) (**Fig. 3A**). In Mg^2+^-limited conditions, bacteria activate dual two-component signaling pathways, PmrAB and PhoPQ, modifying LPS with L-Ara4N and PEtN, leading to resistance to polymyxins(*42–44*) (**Fig. 3A**). Thus, we suspected that fungal-mediated Mg^2+^ depletion might have the unexpected consequence of conferring polymyxin resistance in bacteria.

**Figure 3.**
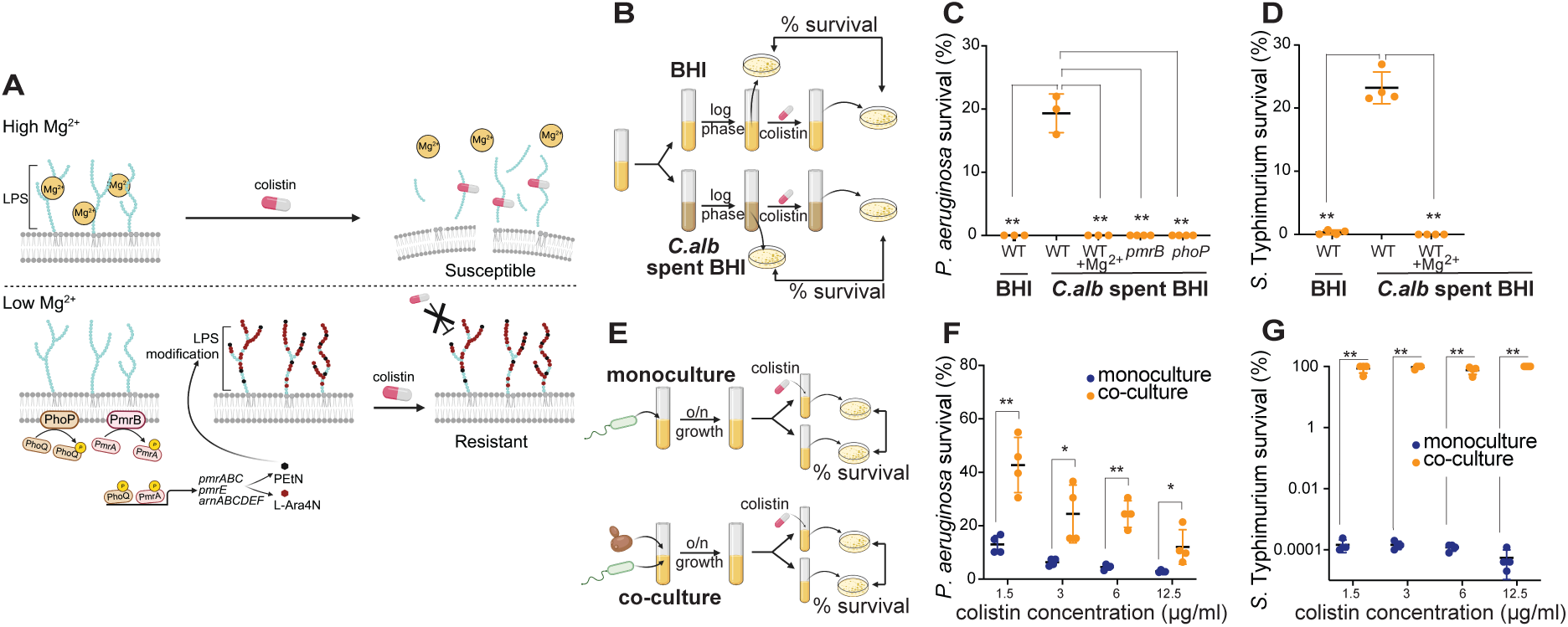
*C. albicans*-mediated Mg^2+^ sequestration protects gram-negative bacteria from colistin. **(A)** Mg^2+^ binds to negatively charged lipopolysaccharides (LPS) on the outer membrane of gram-negative bacteria. In high Mg^2+^ conditions, gram-negative bacteria are sensitive to colistin (and other polymyxin antibiotics), which targets Mg^2+^-bound LPS to disrupt cell membranes, leading to bacterial cell death. Low Mg^2+^ conditions activate two-component signaling pathways, PmrAB and PhoPQ, that modify LPS (PEtN and L-Ara4N), making bacteria resistant to colistin. **(B)** Schematic of the colistin resistance assay. *P. aeruginosa* strains were grown to stationary phase in BHI, *C. albicans* spent BHI, or *C. albicans* spent BHI supplemented with 10 mM Mg^2+^, and then passaged until in exponential phase, when they were treated with 25 μg/ml colistin. Bacterial survival is shown as the percentage of viable cells after the colistin treatment. **(C)** WT *P. aeruginosa* strains were highly susceptible to colistin treatment in BHI media. However, they can survive colistin treatment in *C. albicans* spent BHI media but not in spent BHI supplemented with 10 mM Mg^2+^. In contrast to WT strains, *pmrB* or *phoP* Tn-mutants of *P. aeruginosa* (which cannot modify LPS) remained highly susceptible to colistin even in *C. albicans* spent BHI media. **(D)** WT *S.* Typhimurium was highly susceptible to 3 μg/ml colistin in either BHI media, or in fungal spent media supplemented with 10 mM Mg^2+^, but is more resistant to colistin in fungal spent media alone. **(E)** Schematic of the colistin survival assay. The *P. aeruginosa* WT strain was cultured in BHI only or in co-culture with *C. albicans* for 18 hours. Monocultures or co-cultures were then treated with colistin. Bacterial survival is calculated as the percentage of viable cells after the colistin treatment, just as in (B). **(F)** *P. aeruginosa* co-cultures with *C. albicans* (orange) were more resistant than monocultures (blue) over a series of colistin concentrations, from 1.5 μg/ml colistin (clinical MIC) to 12.5 μg/ml colistin (8x clinical MIC). **(G)** Like in (F), co-cultures of *S.* Typhimurium with *C. albicans* (orange) were significantly more resistant than monocultures of *S.* Typhimurium alone over a range of colistin concentrations. Mean ± std of three biological replicates is shown in panels C-D, F-G. (\*\**p* < 0.01 unpaired two-tailed Student’s *t*-test used).

To test this possibility, we grew *P. aeruginosa* in fresh BHI media or *C. albicans*-spent media, which mimics Mg^2+^ depletion by fungi, and examined bacterial survival when treated with polymyxin E (colistin) (**Fig. 3B**). Consistent with our hypothesis, we found that cells grown in fresh media showed no survival after colistin treatment, whereas cells grown in fungal spent media exhibited a 20% likelihood of survival (**Fig. 3C**). Supplementation of extra Mg^2+^ in fungal-spent media restored the sensitivity of *P. aeruginosa* cells to colistin, just as in BHI fresh media (**Fig. 3C**). As expected, we found that previously characterized *pmrB or phoQ* loss-of-function transposon mutants (*45*) abolished colistin resistance in fungal spent media (**Fig. 3C**). This phenomenon is not restricted to *P. aeruginosa* or colistin. We also found that Mg^2+^ depletion by *C. albicans* increased the survival of *S.* Typhimurium upon colistin treatment, a phenotype which is also suppressed by extra Mg^2+^ (**Fig. 3D**). Further, Mg^2+^ depletion, increased *P. aeruginosa* survival when exposed to another polymyxin antibiotic, polymyxin B (**Fig. S10**). Thus, fungal sequestration of Mg^2+^ provides a general means of conferring polymyxin resistance on gram-negative bacteria.

Fungal spent media cannot fully recapitulate co-culture conditions. Therefore, we tested whether fungal co-culture conditions also confer colistin resistance (**Fig. 3E**). Indeed, we found that transient co-culture with *C. albicans* protects *P. aeruginosa* and *S.* Typhimurium from colistin across a range of concentrations (1.5 μg/mL to 12.5 μg/mL; the clinical MIC of *P. aeruginosa* to colistin is 2 μg/mL) in contrast to monoculture conditions at both 30°C (**Fig. 3F and G**) and 37°C (**Fig. S11A and B**). We also extended our studies to *P. aeruginosa* clinical isolates obtained from patients with cystic fibrosis (CF). Consistently, we found that co-culture with *C. albicans* increased the survival of several colistin-sensitive *P. aeruginosa* CF isolates at a colistin concentration to which these isolates previously displayed sensitivity (**Fig. S12**). Thus, *C. albicans* can protect gram-negative bacteria from colistin across a range of conditions.

### Fungi confer phenotypic, but not genetic, resistance to bacteria against colistin

Understanding how microbial interactions drive the evolution of antibiotic resistance is important for biological and biomedical reasons; increased antibiotic resistance is often observed in polymicrobial communities (*46*). We hypothesized that long-term co-existence with *C. albicans* may fundamentally alter the onset and evolution of colistin resistance in *P. aeruginosa*. To test this possibility, we passaged eight replicate WT populations of *P. aeruginosa* PAO1 for 90 days, either in monoculture or in co-culture with *C. albicans*, with each population exposed to gradually increasing concentrations of colistin (from 1.5 μg/mL to 192 μg/mL) (**Fig. 4A**). As controls, we also passaged mono- and co-culture populations in the absence of colistin (**Fig. S13**). After evolution, both monoculture- and co-culture-evolved populations gained similar resistance to 192 μg/mL colistin (**Fig. 4B**). However, colistin resistance of *P. aeruginosa* from co-culture-evolved populations was dependent on *C. albicans*; colistin resistance was almost completely lost upon removal of *C. albicans* using antifungal treatment (**Fig. 4C**). Our findings indicate that bacteria evolved in co-culture with *C. albicans* acquire non-genetic colistin resistance, due to fungal protection induced by Mg^2+^ sequestration, resulting in continued dependence on the fungus for colistin resistance. This interaction may complicate the treatment of bacterial infections with colistin when *C. albicans* or other fungi are present and suggests a potential benefit to concomitant antifungal therapy.

**Figure 4.**
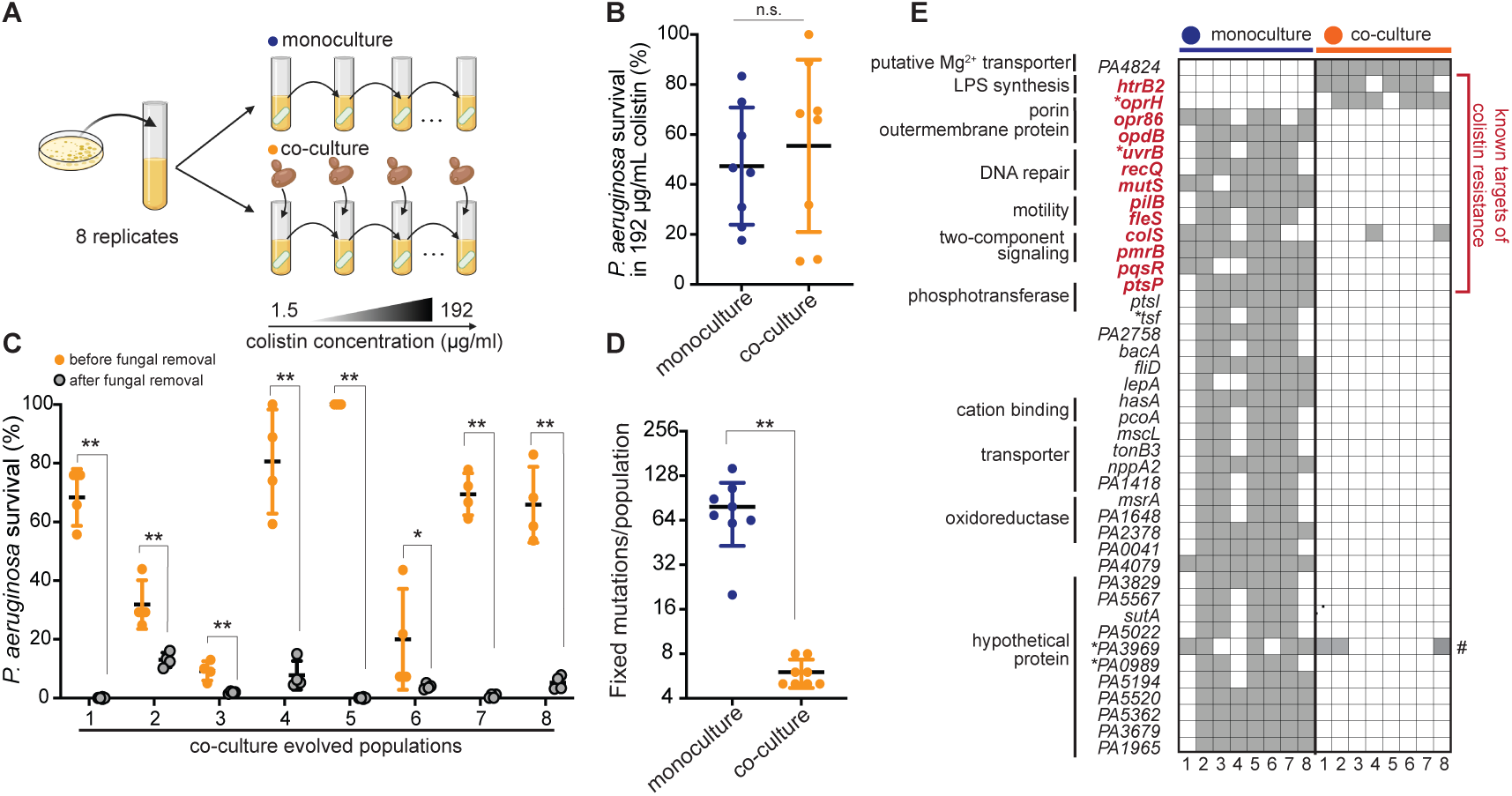
Long-term co-existence with *C. albicans* alters the evolution of *P. aeruginosa* colistin resistance. **(A)** Eight independent WT *P. aeruginosa* populations were passaged daily into BHI with colistin or in BHI with both *C. albicans* and increasing amounts of colistin from 1.5 μg/ml to 192 μg/ml (128x clinical MIC) over a course of 90 days (see Methods). **(B)** After evolution, eight monoculture replicate populations (blue) were not significantly different from eight co-culture populations (orange) for resistance to 192 μg/ml colistin. Co-culture populations were treated with colistin in the presence of co-evolving fungal cells. For each population, the average survival rate of four replicates is shown (n.s. indicates no significance, Mann-Whitney U test). **(C)** *P. aeruginosa* cells from co-cultured-evolved populations lost colistin resistance upon removal of *C. albicans* with antifungal nystatin treatment. Mean ± std of four biological replicates is shown (* *p* < 0.05, ** *p* < 0.01, Mann-Whitney U test). **(D)** Genome sequencing revealed that *P. aeruginosa* strains from either monoculture or co-culture-evolved colistin-resistance populations had different numbers of fixed mutations per population (** *p* < 0.01, Mann-Whitney U test) due to recurrent hypermutator mutations in monoculture populations. **(E)** Monoculture- and co-culture-evolved populations acquire colistin resistance via a distinct spectrum of putative adaptive mutations. Columns indicate each individual replicate monoculture (blue) or co-culture (orange)-evolved population. Rows indicate genes that acquire mutations, grouped by cellular processes. We list all genes with fixed mutations (gray) in five or more replicate populations in either condition (* indicates a mutation in the promoter region). The only genes common between monoculture and co-culture-adapted populations are *colS*, previously known for colistin resistance, and *PA3969* (indicated with a #), which is not unique to colistin-treated populations. A more comprehensive list of all fixed mutations is presented in Table S3 and Table S4.

Based on our findings, *P. aeruginosa* populations evolved in co-culture with fungi likely achieve a phenotypically resistant state but remain genetically susceptible to colistin. Thus, we expect them to have a fundamentally different mutation profile than *P. aeruginosa* cells that have evolved colistin resistance in monoculture conditions. To test this hypothesis, we performed genome sequencing of all experimentally evolved populations. We found that all monoculture-evolved populations became hypermutators (**Fig. 4D**) by acquiring at least one mutation in genes involved in DNA replication or repair (*uvrB*, *recQ*, and *mutS*) (**Fig. 4E and Table S3**); hypermutation is a hallmark of antibiotic-resistance acquisition that has been observed in several previous experimental evolution studies (*47*). As a result of hypermutation, each monoculture population acquired hundreds of fixed mutations (**Fig. 4D**), including mutations in genes known to confer colistin resistance, such as *pmrB* (*48*), *ptsP* (*49, 50*), *pqsR* (*49*), *colS* (*51, 52*), and genes that encode porins (*49*), and protein related to bacterial motility (*53*) (**Fig. 4E and Table S3**). In contrast, co-culture-evolved populations only contained six to eight fixed mutations (**Fig. 4D**), primarily in other targets for colistin resistance, such as genes in LPS biosynthesis (*54*) and an outer-membrane protein *oprH* (*44*) (**Fig. 4E and S14 and Table S3**). Intriguingly, *PA4824*, another co-culture-specific defense gene identified in previous Tn-seq experiments, was also mutated in all eight replicate co-culture-evolved populations (**Fig. 4E and S14 and Table S3**). We did not find fixed mutations in any of these genes in co-culture populations that were not exposed to colistin (**Table S4**). Thus, the dual stressors of colistin and fungal co-culture specifically select for distinct classes of adaptive mutations for colistin resistance.

## Discussion

Nutritional competition for trace metal ions, including iron, zinc, and manganese, either with microbial competitors or hosts, is an important determinant of microbial success during infection. Our study uncovers a novel nutritional competition for the critical Mg^2+^ cation, which fungi sequester from gram-negative bacteria, similar to the Mg^2+^ depletion strategy employed by macrophages to restrict intracellular pathogens (*29*). We show that, under low Mg^2+^ conditions, fungal sequestration of Mg^2+^ may shield bacteria phenotypically from colistin – an antibiotic of last resort against multi-drug resistant *P. aeruginosa* – while at the same time impeding the genetic evolution of colistin resistance. Disrupting this fungal protection by combination therapies involving antifungals may therefore prolong or enhance the efficacy of colistin treatment or might even rescue it even after colistin resistance has been apparently acquired.

Our work thus reveals a new axis of fungal-bacterial competition and suggests rational strategies that may help combat the treatment of multidrug-resistant or seemingly intractable polymicrobial infections.

## Acknowledgements

We thank Dr. Larry Gallagher (Joseph Mougous lab) and Dr. Gina Lewin (Marvin Whiteley lab) for their advice on library construction and data analysis of Tn-seq experiments. We thank Dr. Eduardo Groisman for sharing *S. Typhimurium* strains and Mg^2+^ reporter plasmids. We thank Dr. Colin Manoil and Dr. Pradeep Singh for sharing *P. aeruginosa* strains. We thank Nicole Smalley for sharing her expertise in *P. aeruginosa* genetics. We thank María Angélica Bravo Núñez, E. Peter Greenberg, Carrie Harwood, Meng-Chao Yao, and members of the Dandekar and Malik labs (Ching-Ho Chang, Peter Dietzen, Rechel Geiger, Grant King, Isabel Mejia Natividad, Samantha Wellington Miranda, Maria Toro Moreno), who provided constructive feedback on our study and manuscript. This study was funded by the postdoctoral fellowship from the Cystic Fibrosis Foundation (HSIEH21F0, to YPH), NIH R01 grant GM125714 (to AAD), a Burroughs-Wellcome Fund Career award for medical scientists (1012253, to AAD), and a Howard Hughes Medical Institute Investigator award (to HSM). Funding agencies played no role in study design or publication.

## Author contributions

Conceptualization: YPH, DAH, AAD, HSM Methodology: YPH, WS, JMY Investigation: YPH, WS, RC, JMY Visualization: YPH, WS, AAD, HSM Funding acquisition: YPH, AAD, HSM Supervision: AAD, HSM Writing-original draft: YPH, AAD, HSM Writing-review & editing: WS, JMY, RC, DAH

## Data and materials availability

All data in the main text and supplementary materials are publicly available. Strains and plasmids are available upon request.

## Supplementary Materials

### Materials and methods

#### Bacterial and fungal strains and culture conditions

All bacterial and fungal strains used in this study are listed in **Table S5** in the supplemental material. Bacterial strains in this study were derived from *P. aeruginosa* PAO1, *E. coli* BW25113, and *S.* Typhimurium 14028S. Yeast strains were derived from *C. albicans* SC5314*, S. cerevisiae* w303. *C. tropicalis* CBS94, *C. parapsilosis* CBS604, and *C. glabrata* CBS562 were purchased from ATCC. Individual *P. aeruginosa* Tn mutant were obtained from Dr. Colin Manoil’s lab(*1*); correct transposon insertions were confirmed by PCR and Sanger sequencing. Clinical *P. aeruginosa* isolates from CF patients were obtained from Dr. Pradeep Singh’s lab. Unless otherwise specified, all experiments were performed in Brain Heart Infusion Broth Media (BHI, Sigma-Aldrich), buffered with 10% MOPS to pH7.0 followed by filter sterilization. All strains were grown in BHI and incubated at 30°C or 37°C. Other media used in this study were YPD (1% yeast extract, 2% peptone, and 2% D-glucose), Synthetic Cystic Fibrosis Medium (SCFM(*2*)), and Tryptic Soy Broth (TSB, Sigma-Aldrich). For antibiotics used in this study, 100 μg/mL gentamicin was used to select against Gram-negative bacteria. 50 μg/mL nystatin was used to select against *Candida* species and *S. cerevisiae*. Colistin and polymyxin B (Sigma-Aldrich) were prepared as 10 mg/mL and 5 mg/mL stock solutions respectively.

#### Generation of *P. aeruginosa* mutants

For generating gene deletions in *P. aeruginosa*, pEXG2-mediated allelic exchange method(*3*) was used. In short, pEXG2 deletion constructs were transformed into *E. coli* S17. S17 donor was subsequently mixed with *P. aeruginosa* recipients on an LB agar plate at a 5:1 ratio of donor to recipient cells, and the cell mixture was incubated at 30°C overnight. The cell mixture was then scraped and resuspended into 200μl LB, and plated on LB agar plates with 100 μg/mL gentamicin to select for cells containing the deletion plasmid integrated into the *P. aeruginosa* genome. *P. aeruginosa* merodiploid colonies were streaked on LB-no salt agar plates with sucrose. Gentamicin-sensitive and sucrose-resistant colonies were screened for allelic replacement by colony PCR, and gene deletions were confirmed by Sanger sequencing of PCR products. Primers and Plasmids used for strain constructions are listed in **Table S6**.

#### Generation of *S. cerevisiae* mutants

*S. cerevisiae* gene deletion mutants were generated by homologous recombination at endogenous gene loci. Gene deletion fragments were amplified from pFA6a-natMX4 and fused with 500 bp upstream and downstream sequences of the targeted coding sequence by PCR. The PCR fragment with homology arms was transformed into *S. cerevisiae* cells with the standard LiOAc-based protocol(*4*). Gene deletions were confirmed by colony PCR and Sanger sequencing to check the absence of endogenous genes. Primers and Plasmids used for strain constructions are listed in **Table S6**.

#### Fungal-bacterial co-culture CFU assay

All experiments began with bacterial or fungal cultures that were grown overnight. The starter cultures were diluted either 1:100 (for bacterial cultures) or 1:50 (for fungal cultures) in BHI and cultured for ∼4-5 hours to reach log phase. Refreshed bacterial and fungal strains were added to BHI to reach final bacterial cell density at 2.5 x 10^4^ cells/mL and fungal cell density at 5 x 10^5^ cells/mL, as the co-culture experiment. The same amounts of bacterial or fungal cells were also added separately in fresh BHI, as monoculture controls. These cultures were incubated for 18 hours at 30°C or 37°C with shaking. Cultures were serially diluted and spotted on LB plates with 50 μg/mL nystatin for measuring bacterial growth and on YPD plates with 100 μg/mL gentamicin for measuring fungal growth.

#### Measurements of bacterial intracellular Mg^2+^

Cytosolic Mg^2+^ concentrations in *P. aeruginosa* were measured using a modification of the previously reported Mg^2+^ genetic sensor assay(*5*). First, we cloned the Mg^2+^ genetic sensor containing *S. Typhimurium mgtA* leader sequence (*P_lac_*-*mgtA_leader_-lacZ*) or the promoter-only fragment (*P_lac_*-*lacZ*) into a *P. aeruginosa* replicating plasmid, pBBR. Then, these two plasmids were transformed into the indicated *P. aeruginosa* strains separately, with the former acting as a reporter for Mg^2+^ and the latter acting as a control. Both strains were cultured in BHI or fungal spent media (80% v/v final concentration) containing 100 μg/mL gentamicin for 16 hours and then diluted 1:100 in the corresponding media with gentamicin for 4 hours to reach log phase. The cell density of these cultures was adjusted to OD600=0.4, and we followed the same protocol to measure β-galactosidase activity using Galacto-Light Plus System (ThermoFisher). Finally, cytosolic Mg^2+^ concentrations were derived from the following equation, where the control strain was used to normalize transcription or translation efficiency between strains: [Mg^2+^] = (miller units from the control strain)/ (miller units from the reporter strain)

#### Colistin resistance assay

To prepare *C. albicans*-free supernatant, a single *C. albicans* colony was inoculated in 10mL BHI and cultured at 30°C for 24 hours. The fungal culture was centrifuged, and the supernatant was filtered with a Steriflip unit with a 0.22 μm Millipore Express PLUS membrane (MilliporeSigma). 8mL fungal filtrate was added to 2 mL fresh BHI (80% v/v final concentration) to prepare fungal spent media, mimicking nutrient exhaustion by *C. albicans*. To conduct colistin treatment, overnight bacterial culture was diluted in 3 mL BHI or fungal spent media at 1:100 for 4 hours. Then, bacterial cultures were adjusted into OD600=0.3 in BHI or fungal spent media. Each bacterial culture was treated with colistin at the indicated concentration, and the samples were incubated at 30°C for 1.5 hours. To assay bacterial viability, bacterial cultures after colistin treatment were serially diluted on LB plates. Bacterial cultures in the same growth condition without colistin treatment were used as a control. Bacterial survival was calculated by the ratio of bacterial CFUs with, relative to without treatment, at 1.5 hours.

#### Colistin survival assay

Log-phase bacterial cells were cultured in BHI alone or co-culture with fungal cells in BHI for 18 hours at 30°C or 37°C, as the co-culture CFU assay indicated. To equilibrate bacterial cell numbers in these two culturing conditions prior to the colistin treatment, the cell density of monoculture samples was adjusted to OD600=0.3 to match the bacterial cell density after 18-hour growth in co-culture. Both equilibrated monoculture and co-culture samples were split into 1mL in two different tubes for colistin-treated versus untreated conditions. Colistin was added to the first tube at the indicated concentration. Bacterial cells in both tubes were incubated at 30°C for 1.5 hours and the bacterial viability was determined by counting CFUs after serial dilution. Bacterial survival was determined by the ratio of bacterial CFUs upon colistin treatment, relative to without treatment (average of four replicates). Bacterial survival was set to 100% if it was over 100% due to stochastic variation in colony counts.

We used the same fungal protection assay to measure colistin resistance in co-culture-evolved populations. These evolved populations were split into two culturing conditions: BHI only, as co-culture conditions, or BHI + 50 μg/mL nystatin, to remove fungal cells (as monoculture conditions). Cells were grown for 18 hours at 37°C. After overnight growth, populations in both conditions were treated with 192 μg/mL colistin following the above protocol. To measure bacterial viability after colistin treatments, cells in co-culture and monoculture conditions were spotted on LB + 50 μg/mL nystatin or LB plates, respectively.

#### Fitness competition assay

A strain PAO1 WT with chromosomally integrated miniTn7-mCherry was used as the reference strain in fitness competition with deletion mutants, either in BHI only or in co-culture with *C. albicans*. All bacterial and yeast strains were incubated in BHI at 30°C to log phase, and cell density was quantified by OD600 using a spectrophotometer (biowave CO8000 Cell Density Meter). First, a non-fluorescent strain, either the PAO1 wild type or deletion mutants, was mixed with the fluorescence-labeled reference strain at a ratio of 1:1 to make a bacterial mixed culture with 2×10^6^ cells/mL. Part of the cell mixture was used to determine the initial ratio of sample and reference strains using flow cytometry (BD FACSymphony A5 Cell Analyzer). Then, 10μl of cell mixture was inoculated separately in 3mL BHI alone (monoculture) or 3mL BHI with 2×10^5^ *C. albicans* cells (co-culture) and grown in BHI at 30°C for 18 hours. After 18 hours, monoculture samples were diluted 100-fold to measure the ratio of sample and reference strain. For co-culture samples, a low-speed spin (1000x*g* for 3min) was applied to separate bacterial and fungal populations. Supernatants enriched with bacterial cells were measured using flow cytometry. For each sample, at least 30,000 cells were collected and the FACS data were analyzed by the FlowJo 10.4.1 software. The reference strain was cultured separately to estimate the number of generations during the experiment. Each experiment was conducted in at least two biological and two technical replicates. To calculate the relative fitness, *w*, of each sample strain to the reference strain, we followed the formula: *w*= 1+*s*, where selection coefficient s = [ln(sample/reference)_t_-in(sample/reference)_t=0_]/t, where t= number of generations and (sample/reference) is the ratio between a sample strain and the reference strain(*6*).

#### RNA sequencing and analysis

A log-phase *P. aeruginosa* WT culture (OD600=0.1∼0.2) was diluted at 5×10^6^ cells in 100mL media alone or 100mL media with 5×10^7^ *C. albicans* cells and grown at 30°C. Cells were collected for extracting RNA at 8 hours when there was no significant growth difference of *P. aeruginosa* between monoculture and co-culture. Prior to RNA extraction, a slow-speed spin was applied to co-culture samples to enrich bacteria in the supernatant. For each sample, total RNA was harvested from 10^9^ cells, preserved in RNA-protected QIAzol, and purified by RNAeasy mini kit (Qiagen) following the commercial protocol. RNA was quantified by Qubit fluorometric quantitation system (ThermoFisher), and RNA integrity was examined by 4200 TapeStation System (Agilent). We generated an rRNA depleted library, and obtained > 20 million 150 bp pair-end Illumina NovaSeq reads \ for each sample commercially (Azenta). Reads were examined using FastQC (Barbaham Bioinformatics, Babraham Institute) and trimmed by Trim Galore! (v0.6.10; https://github.com/FelixKrueger/TrimGalore). Reads were mapped to the *P. aeruginosa* PAO1 genome (GCF_00006765.1) using Subread/ FeatureCounts (https://github.com/ShiLab-Bioinformatics/subread). Differential gene expression was analyzed with DESeq2(*7*) (https://bioconductor.org/packages/release/bioc/htmL/DESeq2.htmL), following the default pipeline of data normalization. A log_2_ fold change cutoff of 2 and a false discovery rate (FDR) of 0.05 were used to filter significantly differentially expressed genes between samples from three biological replicates. Gene Ontology analysis was done using the *P. aeruginosa* STRING database on ShinyGO 0.77(http://bioinformatics.sdstate.edu/go/). Sequencing files are stored in the BioProject PRJNA1021673 of the NCBI repository.

#### Transposon-sequencing and analysis

We used a pool of ∼10^5^ transposon-insertion mutants, generated by Tn5-based transposon T8 (IS*lacZ*hah-tc) insertion in the *P. aeruginosa* strain PAO1(*8*). 20μl of frozen Tn mutants (approximately containing 10^8^ cells) were thawed and inoculated in 50mL media or 50mL media with 10^8^ *C. albicans* cells in a 250mL flask, to establish monoculture and co-culture conditions, each with three biological replicates. To screen the fitness of Tn mutants effectively, cells were grown at 30°C with shaking at 210rpm for 10 hours (approximately 10 generations) and collected for genomic DNA extraction using the DNAeasy Blood& Tissue kit (Qiagen). DNA samples were prepared for transposon sequencing following the previously published TdT/ two-step PCR method(*9*). In short, 2μg DNA was end-repaired by NEB end-repair reaction and subsequently C-tailed in the TdT tailing reaction. Two PCR reactions were used to enrich DNA fragments spanning the junction of transposon insertion. PCR-1 was performed using a transposon-specific primer and a polyG primer (olj376) on 0.5-1μg DNA; at least 2 reactions which were pooled in the following purification steps. For PCR-2, a mixture of Tn8 primers combined with 3N, 5N, 7N, and 9N random bases were used to increase the diversity of Tn-seq libraries. The size of library DNA was examined by 4200 High Sensitivity DNA TapeStation System (Agilent) and quantified using qPCR with standard Illumina primers (Kapa Biosystems). Libraries were pooled and sequenced commercially using an Illumina NovaSeq or NextSeq sequencer (Novogene). For each sample, at least 25 million pair-end 150bp sequencing reads were generated after which, we proceeded for downstream analysis. Sequencing files are stored in the BioProject PRJNA1021673 of the NCBI repository.

The analysis scripts used in this study are adapted from previously published protocols and are available at https://github.com/PhoebeHsieh-yuying/P.-aeruginosa_Tnseq_paper. Briefly, sequencing reads were groomed by FastQC (Barbaham Bioinformatics, Babraham Institute), filtered to keep those with the correct Tn8 sequence (TATAAGAGTCAG) near the 5’ end using Cutadapt(*10*), and mapped to the *P. aeruginosa* PAO1 genome (GCF_00006765.1) with Bowtie2(*11*) using the Tnseq2.sh script. The output table with transposon insertion site assignments of each sample was concatenated and annotated with known gene features of *P. aeruginosa* using a custom R script, Tnseq_reformat.R. Then, DESeq2(*7*) statistical package in R was used to identify genes with significant changes in Tn insertion between co-culture and monoculture. A log_2_ fold change cutoff of 1 and a false discovery rate (FDR) of 0.05 were used as the criteria for significance.

#### Experimental evolution of colistin resistance

Eight independent colonies of a WT *P. aeruginosa* strain PAO1 were used as ancestral strains for the experimental evolution experiment. Each ancestral clone was inoculated and grown in BHI at 37°C to 10^9^ cells/mL. At the beginning of the experiment, approximately 10^7^ cells of each ancestral population were transferred either into 1 mL BHI with colistin or into 1mL BHI with colistin and 10^6^ *C. albicans* cells to select colistin resistance. Each day, populations were passaged at a ratio of 1:200 in the former condition, while co-cultured populations were passaged at 1:50 in colistin-containing BHI to maintain comparable population size between both conditions. The concentration of colistin was started from 1.5 μg/mL, gradually increased two-fold every week or every two weeks and reached 192 μg/mL at the end of the 90-day evolution experiment. As controls for adaptation to BHI only or co-culture with *C. albicans*, the same ancestral populations were passaged at a ratio of 1:200 into 1 mL BHI or 1 mL BHI with the same amount of *C. albicans* cells, but without colistin. After every 7 days, 1 mL evolving cultures were mixed with 500 μl 80% glycerol and frozen at −80°C. On day 90, we expected that populations evolved in colistin-containing BHI with *C. albicans* had been passaged for ∼600 generations, while the rest of the evolved populations had been passaged for ∼700 generations.

#### Whole Genome Sequencing and analysis

We sequenced two ancestral populations and all eight evolved populations per treatment. We revived each population from a freezer stock in the growth condition under which they evolved and grew for 24 hours. For co-culture evolved populations, we treated them with nystatin to remove fungal cells. 3 mL bacterial culture was collected for DNA extraction using the DNAeasy Blood&Tissue kit (Qiagen). Sequencing libraries were made and sequenced commercially by Illumina sequencers in SeqCenter (https://www.seqcenter.com/). Variant calling was done using the breseq(*12*) software v0.37.1 and the *P. aeruginosa* PAO1 genome (GCF_00006765.1) was used as the reference genome. The average depth of sequencing for populations was 222.5 fold (+/- 32.7) and the average genome coverage was 98.8 fold (+/- 0.04). Variants at >95% in each population were filtered for mutation calling. Mutations were manually curated by identifying unique variants in each evolved population compared to the ancestor. Sequencing files are stored in the BioProject PRJNA1021673 of the NCBI repository.

#### Mg^2+^ measurements in media

Mg^2+^ concentrations were measured using the Magnesium Assay kit (Sigma-Aldrich) according to the protocol provided. Absorbances at OD_450_ after the enzymatic reaction were compared to a standard curve of Mg^2+^ concentration to determine the absolute concentration in media.

#### Figure Preparation

Cartoon diagrams in this study were prepared by BioRender (https://www.biorender.com/)

## Supplementary Figures

**Figure S1.**
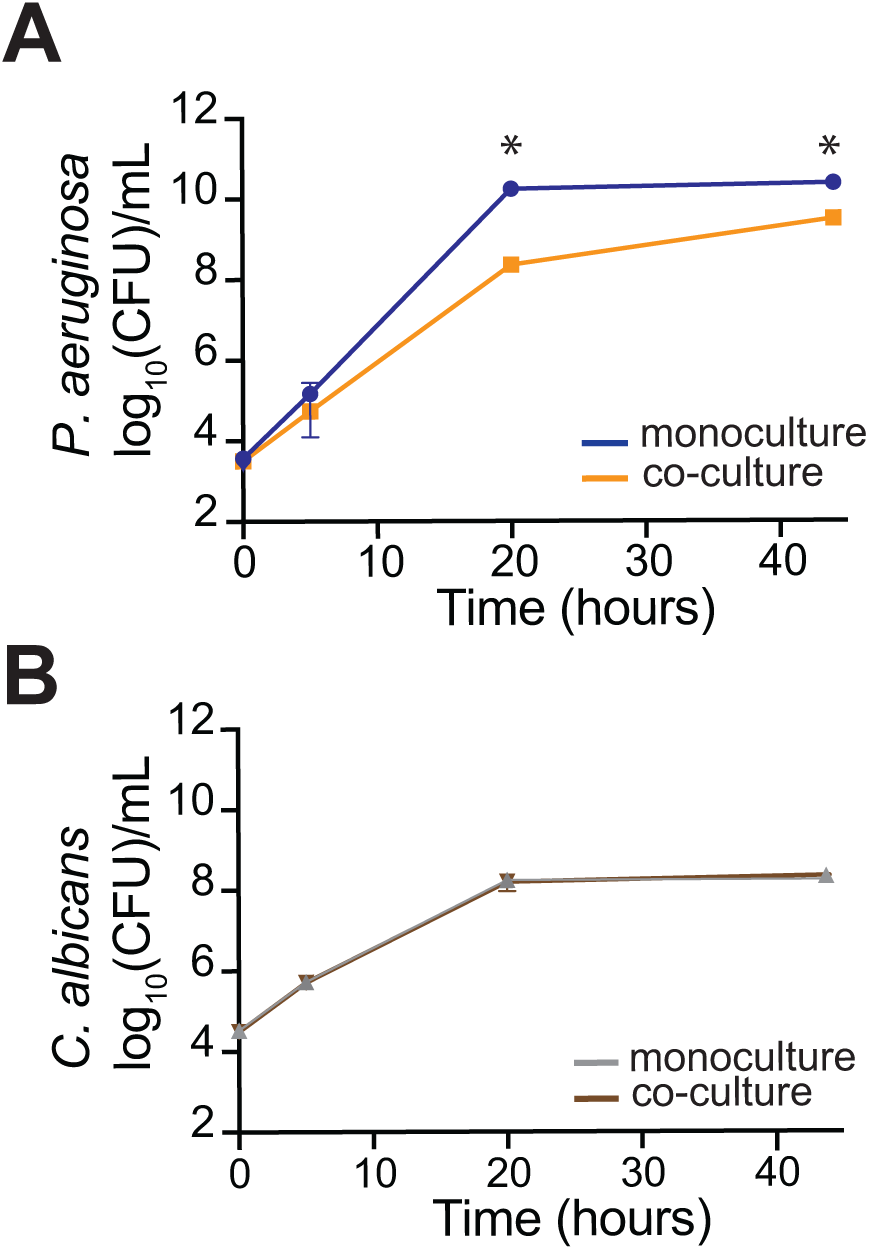
*P. aeruginosa* fitness is suppressed by *C. albicans* in BHI. During co-culture in BHI for 42 hours, *P. aeruginosa* fitness was significantly impaired relative to monoculture, while *C. albicans* fitness remained unchanged (See Methods). The fitness of *C. albicans* or *P. aeruginosa* is shown as colony formation units (CFUs) per mL in (A) or (B), respectively. Mean ± std of three biological replicates is shown (* *p* < 0.05, unpaired two-tailed Student’s *t*-test used).

**Figure S2.**
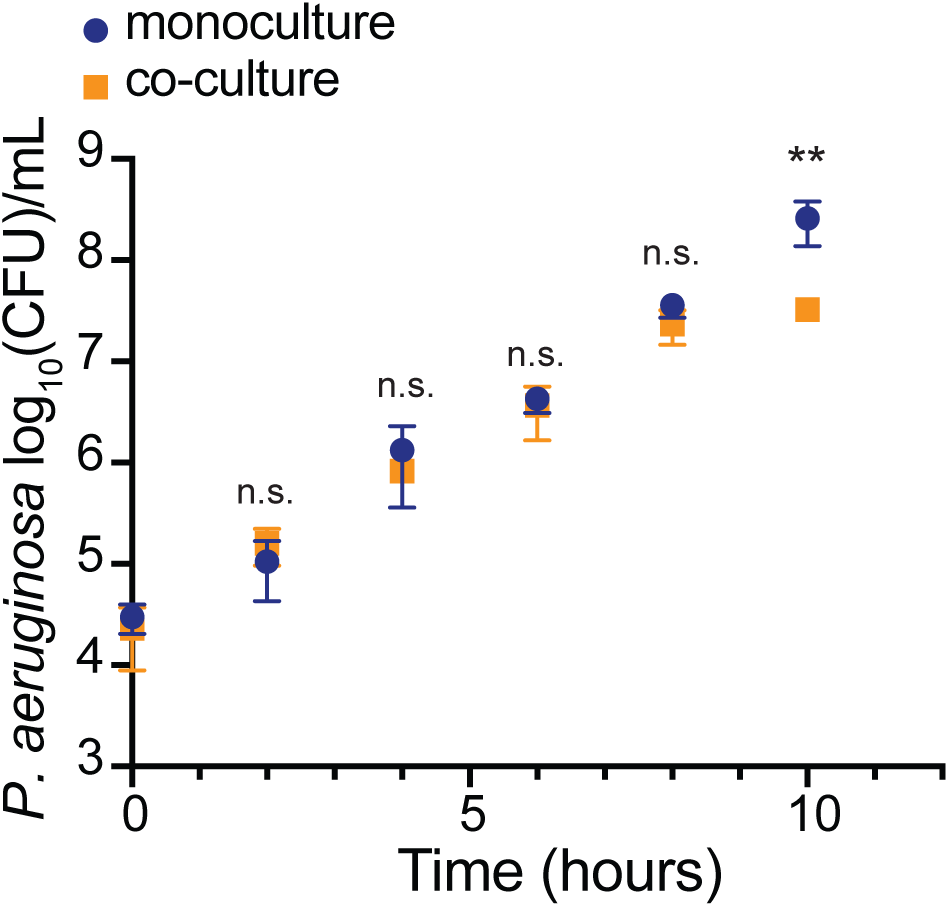
The fitness of *P. aeruginosa* remains unchanged during the first eight hours in co-culture with *C. albicans*. The WT *P. aeruginosa* strain was grown in BHI media only or in co-culture with *C. albicans* (See Methods). The fitness of *P. aeruginosa* was examined every two hours in BHI (blue) and co-culture (orange). Mean ± std of three biological replicates is shown (** *p* < 0.01 and n.s. indicates no significance. Unpaired two-tailed Student’s *t* test was used, relative to monoculture at each time point).

**Figure S3.**
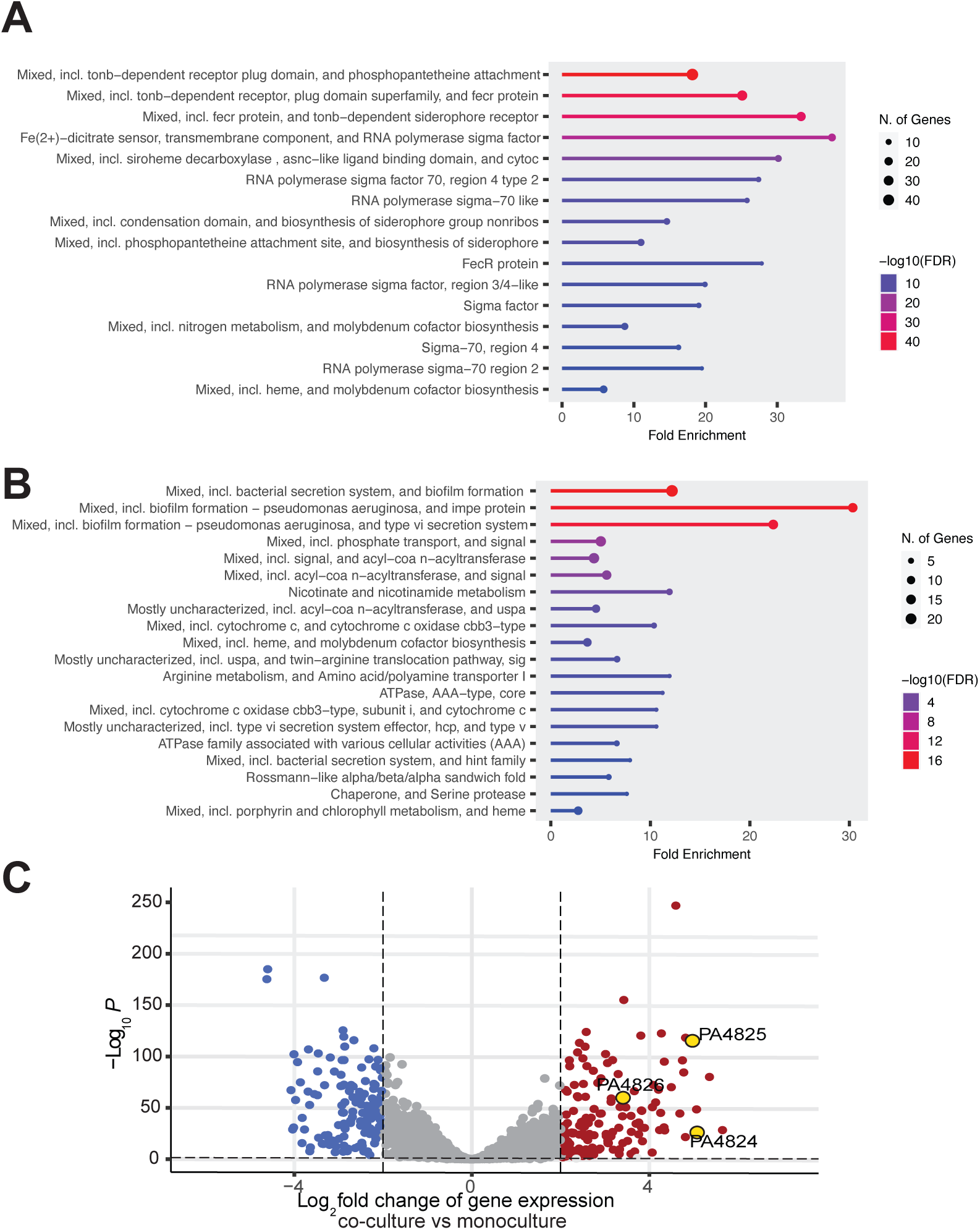
Differentially expressed *P. aeruginosa* genes in co-culture relative to monoculture conditions. Gene ontology analysis of genes up-regulated or down-regulated in co-culture compared to monoculture is shown in (A) or (B), respectively. Up-regulated genes were enriched for iron uptake and RNA polymerase sigma factor 70, while down-regulated genes were enriched for bacterial secretion, co-factor biosynthesis, and energy generation. X-axis indicates fold enrichment of genes of interest, and Y-axis indicates GO categories, which are ordered and color-coded by the inverse of their false discovery rate. The number of genes of interest in each category is labeled with the size of circles. (C) RNA-seq volcano plot (See Methods) shows *P. aeruginosa* genes that were differentially expressed in co-culture versus monoculture conditions. X-axis indicates fold change, while Y-axis indicates *p*-value after correcting multiple tests. |log_2_ fold change| > 2 and adjusted *p* < 0.05 was used as statistical cut-offs. Up-regulated genes are labeled in red, while down-regulated genes are in blue.

**Figure S4.**
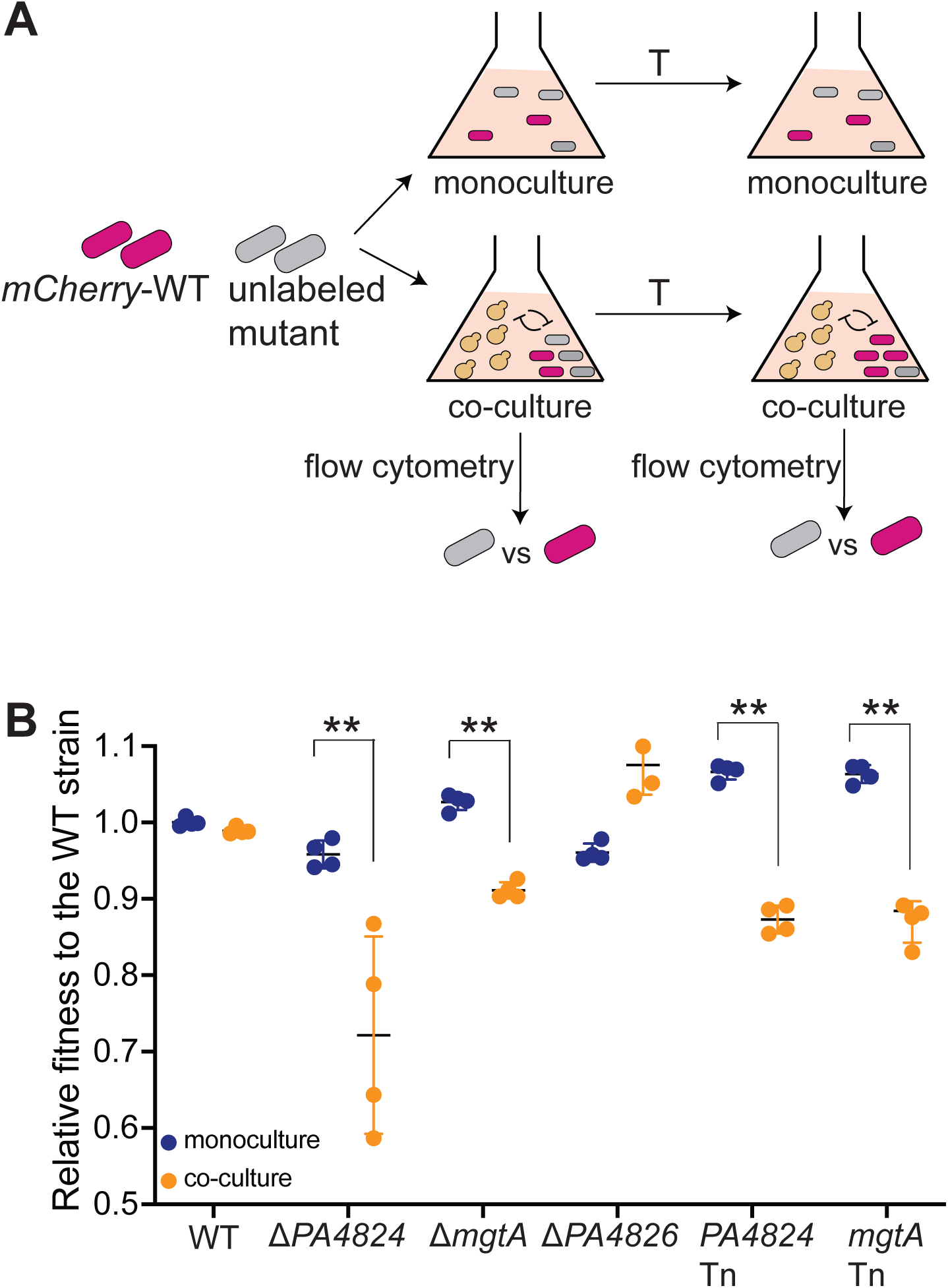
Fitness competition assay verifies the fitness effect of *PA4824* or *mgtA* mutants in co-culture. (A) Schematic of the fluorescence-based fitness competition assay that mimics bulk selection in Tn-seq (See Methods). (B) Relative fitness of *PA4824* or *mgtA* mutants to the WT *P. aeruginosa* strain was impaired in co-culture conditions. Relative fitness in BHI media only is indicated in blue, and relative fitness in co-culture with *C. albicans* is indicated in orange. Mean ± std of three biological replicates is shown (** *p*< 0.01, unpaired two-tailed Student’s *t*-test used).

**Figure S5.**
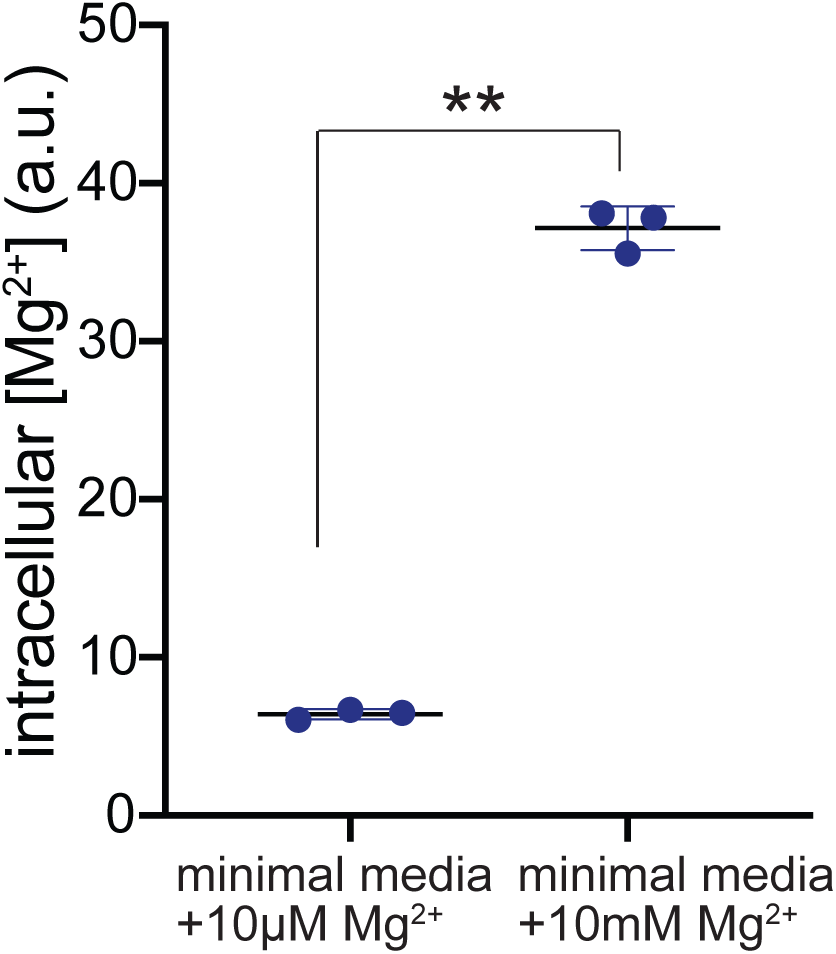
Verification of the Mg^2+^ genetic sensor in *P. aeruginosa*. The WT *P. aeruginosa* strain carrying the Mg^2+^ genetic sensor (pBBR-P*_lac1-6_-mgtA_leader_-lacZ*) was cultured in minimal medium with 10μM or 10mM Mg^2+^ to log phase, mimicking low or high Mg^2+^ conditions. Relative concentrations of intracellular Mg^2+^ were determined by the measurements of β-galactosidase activity (See Methods). Mean± std of three biological replicates is shown (** *p* < 0.01, unpaired two-tailed Student’s *t*-test used).

**Figure S6.**
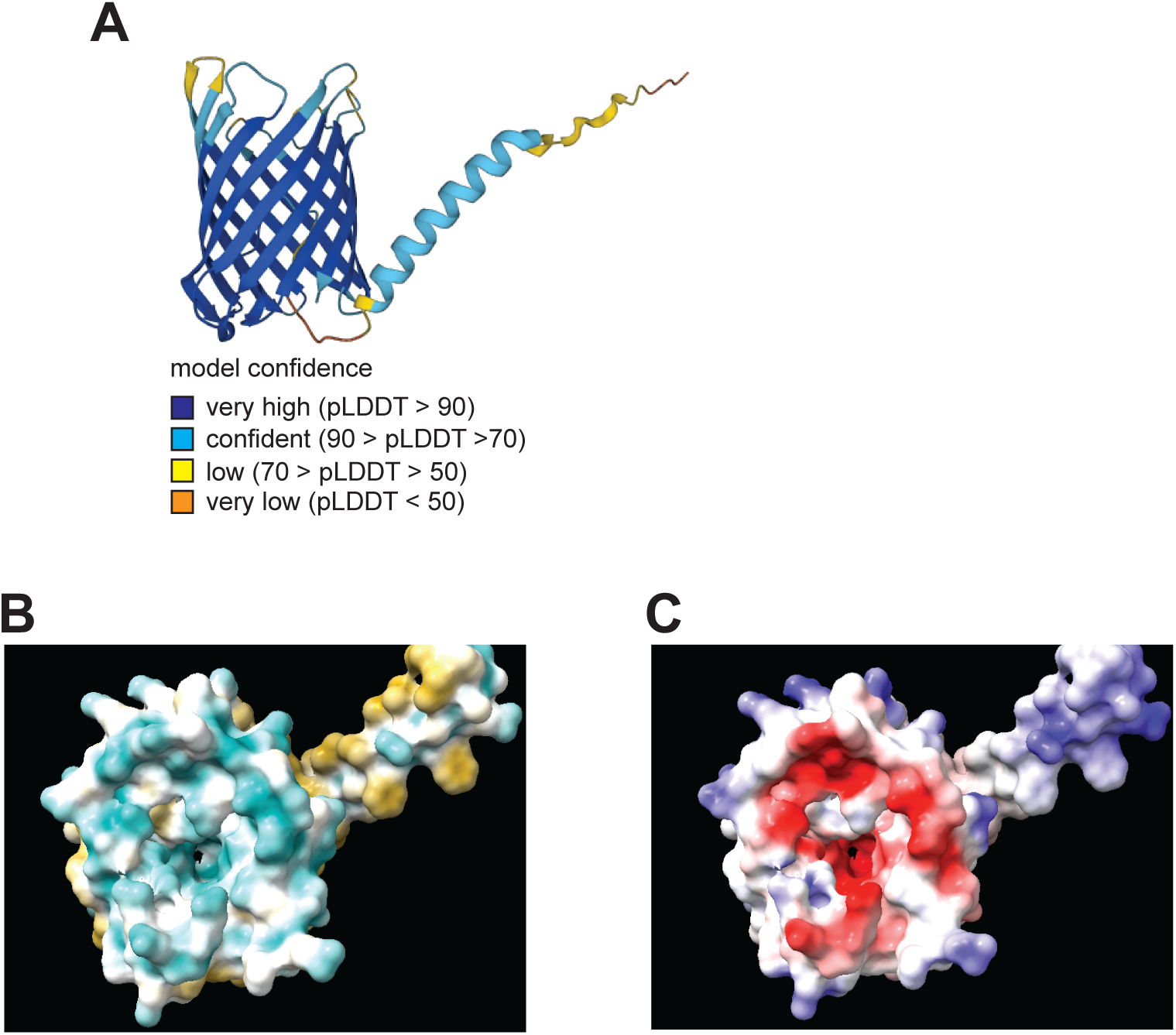
Alphafold structural prediction suggests PA4824 may function as a Mg^2+^ transporter. (A) An Alphafold-predicted structure of PA4824. Confidence in structural prediction is shown by color, where dark and light blue indicate trustable confidence. (B) and (C) show the top-down view of the core of PA4824, displayed by Chimerax. In (B), residues are labeled by their hydrophobicity (green: hydrophilic residues, yellow: hydrophobic residues). In (C), residues are labeled by their electrostatic features (red: negatively-charged residues, blue: positively-charged residues).

**Figure S7.**
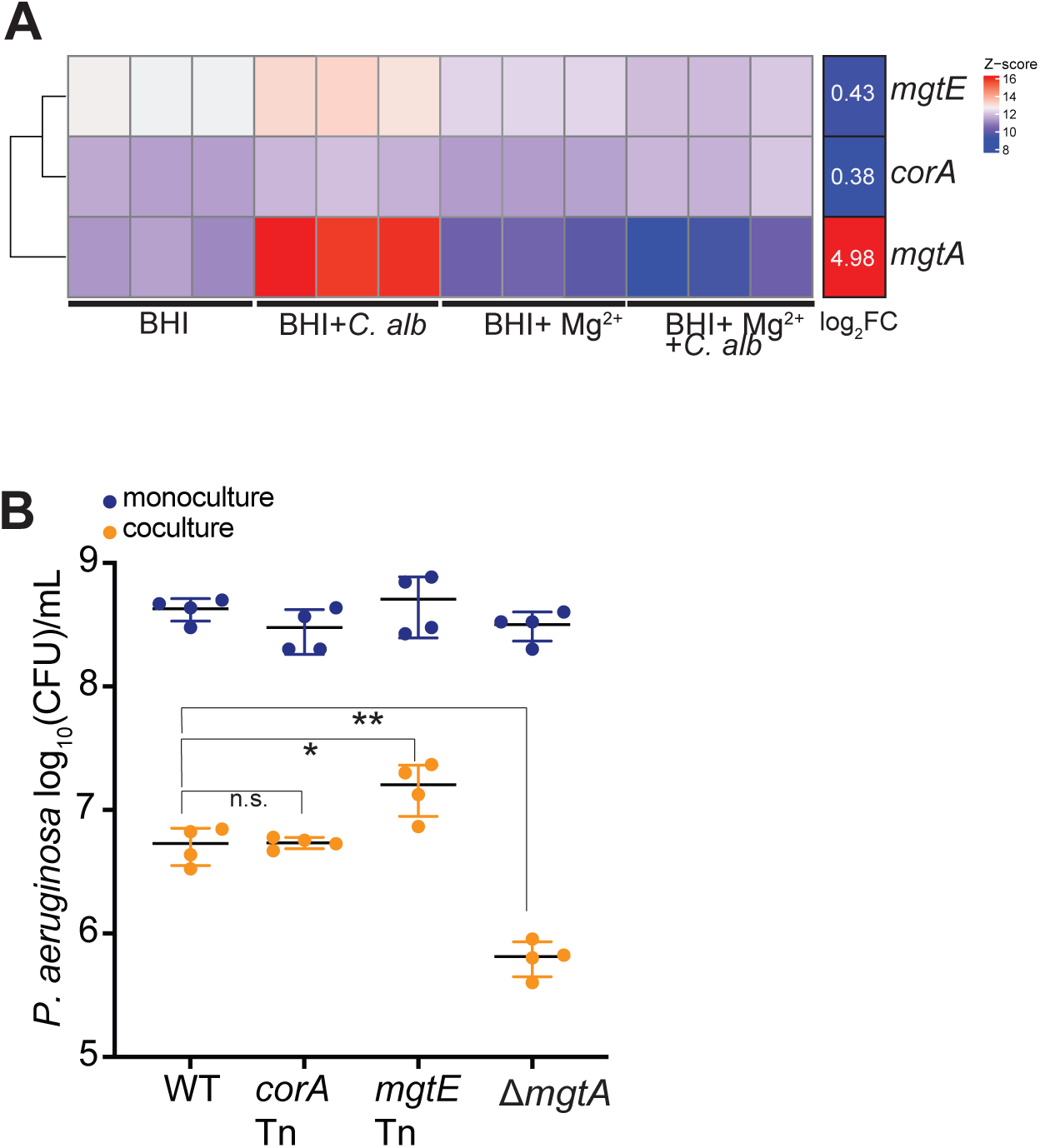
Two other *P. aeruginosa* Mg^2+^ transporters, CorA and MgtE, are not involved in Mg^2+^ competition with *C. albicans*. (A) Expression levels of *mgtE* and *corA* were not changed in responsive to the presence of *C. albicans*. RNA-seq results of *mgtE*, *corA*, and *mgtA* across four experimental conditions (BHI only, co-culture in BHI, BHI with supplemental Mg^2+^ (10mM), and co-culture in BHI supplemental Mg^2+^ (10mM)) are shown in z-score. The log_2_ fold change of each gene in co-culture versus monoculture in BHI is shown on the right. (B) *corA* or *mgtE* Tn mutants did not confer fitness cost in co-culture with *C. albicans*, as Δ*mgtA* mutant did. Fitness of *P. aeruginosa* strains in BHI only or co-culture conditions are shown in blue or orange, respectively. Mean ± std of four biological replicates is shown (** *p* < 0.01, * *p* < 0.05, and n.s. indicates no significance. Unpaired two-tailed Student’s *t*-test was used)

**Figure S8.**
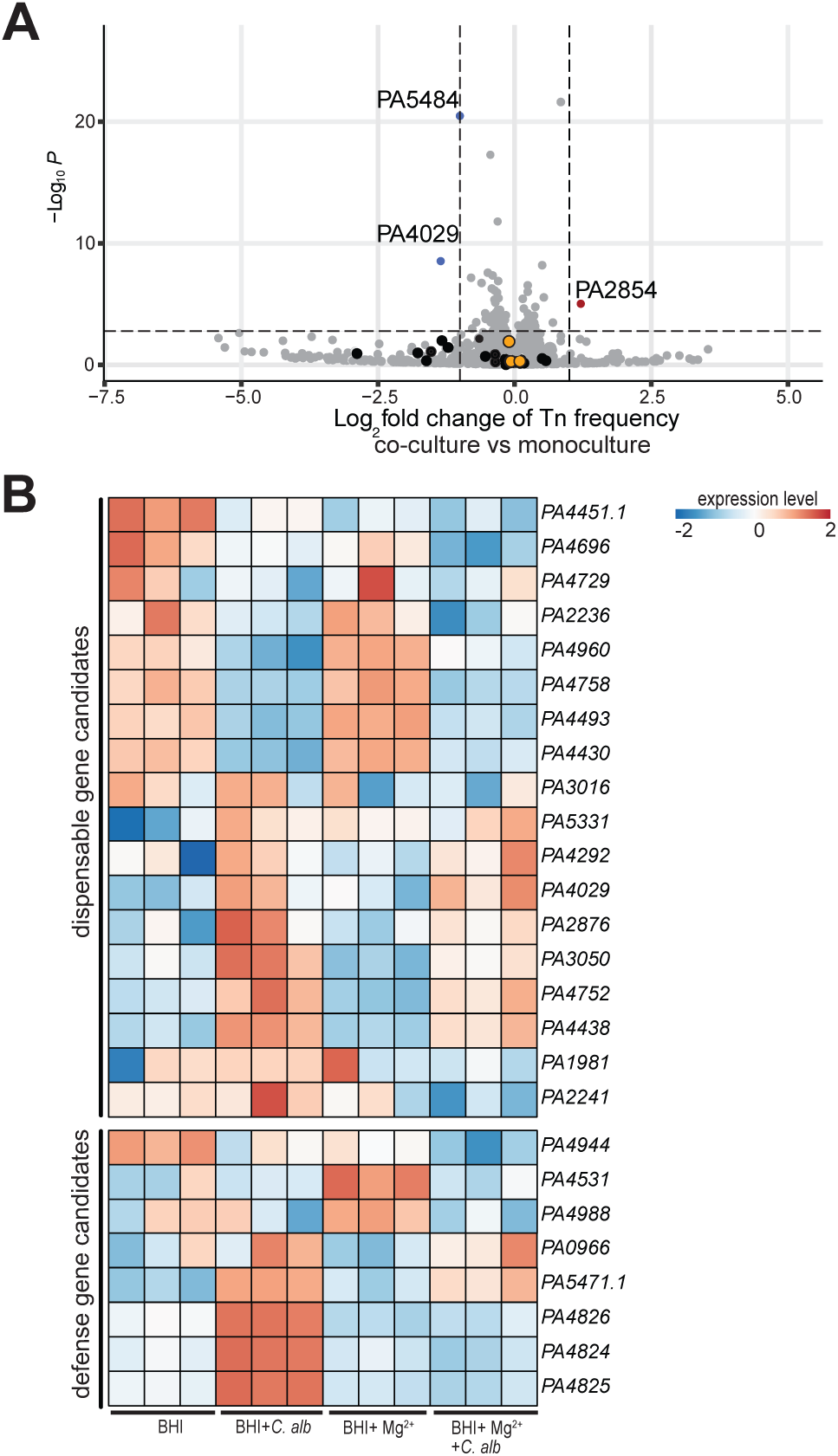
Mg^2+^ supplementation rescues the fitness effects of other candidate genes, but does not change their expression. (A) Tn-seq volcano plot in Mg^2+^-repleted conditions (See Methods) shows the fitness effect of most of the candidate defense and dispensable genes (in black) were restored. X-axis indicates fold change, while Y-axis indicates p-value after correcting for multiple testing. We used log_2_ fold change > 2 and adjusted *p*-value < 0.1 as statistical cut-offs. *PA4824*, *PA4825*, and *PA4826* are labeled in yellow. Under this condition, only one gene was dispensable (in red, *PA2854*), and two genes were essential (in blue, *PA5484* and *PA4029*). (B) Mg^2+^ supplementation only altered the expression of *PA4824*, *PA4825*, and *PA4826* in co-culture, but not the other candidate genes. Heatmap shows the expression of 18 dispensable genes and eight defense genes in co-culture versus monoculture, with or without Mg^2+^ supplementation.

**Figure S9.**
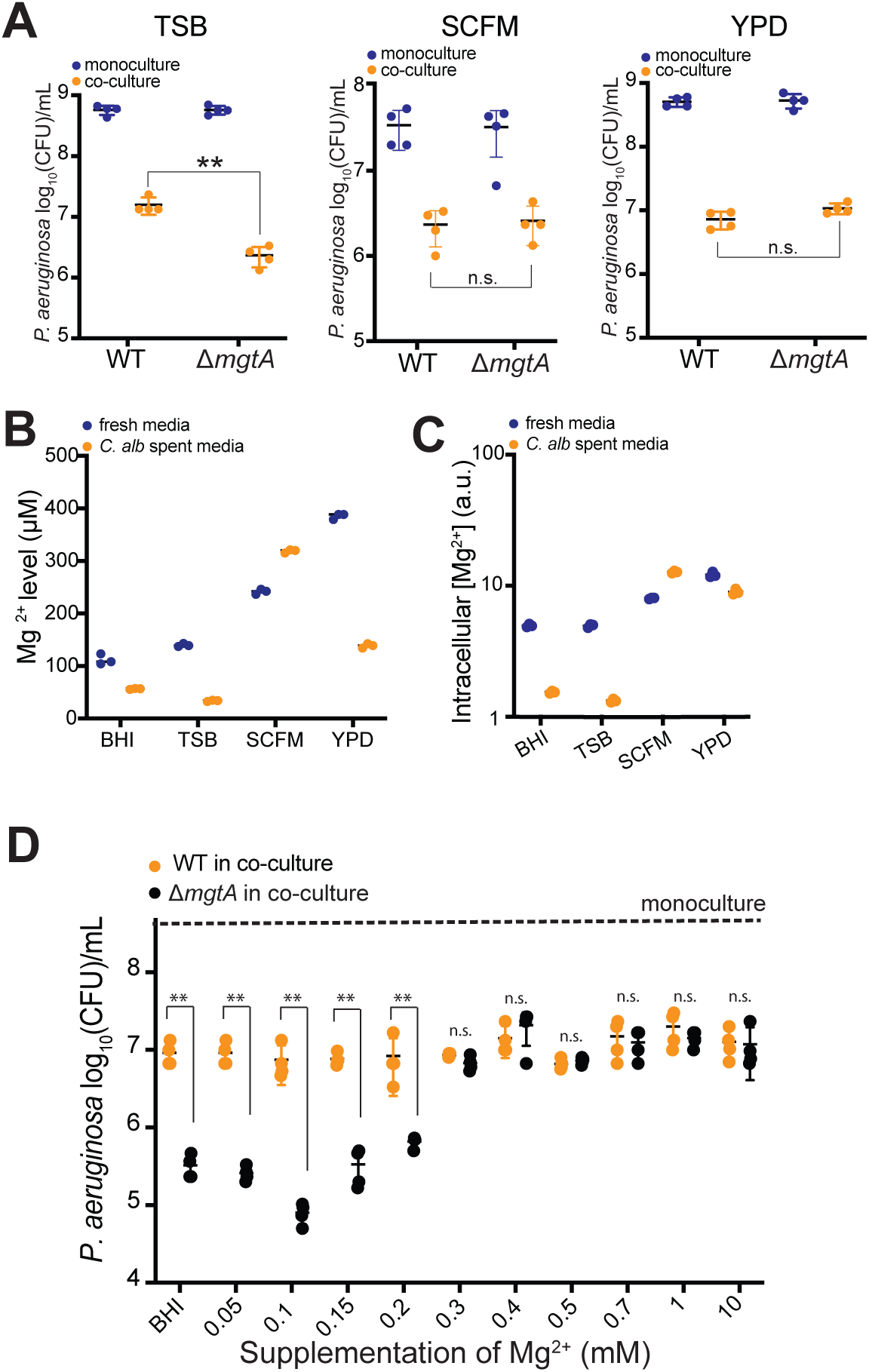
Mg^2+^-sequestration by fungi depends on the environmental Mg^2+^ level. (A) Fitness of *P. aeruginosa* Δ*mgtA* mutant was also impaired in co-culture with *C. albicans* in TSB, but not SCFM, and YPD. Bacterial fitness in media only or in co-culture is shown in blue or orange, respectively. Mean ± std of four biological replicates is shown (** *p* < 0.01 and n.s. indicates no significance. Unpaired two-tailed Student’s *t*-test was used) (B) BHI and TSB exhibited lower Mg^2+^ levels compared to YPD and SCFM, both in the fresh media or *C. albicans* spent media. Mean ± std of three biological replicates is shown. (C) Intracellular Mg^2+^ levels in *P. aeruginosa* were lower in the fungal spent media from BHI and TSB. Mean ± std of four biological replicates is shown. (D) Co-culture-specific fitness loss of the *P. aeruginosa* Δ*mgtA* mutant was rescued by adding at least 0.3 mM Mg^2+^ in BHI.

**Figure S10.**
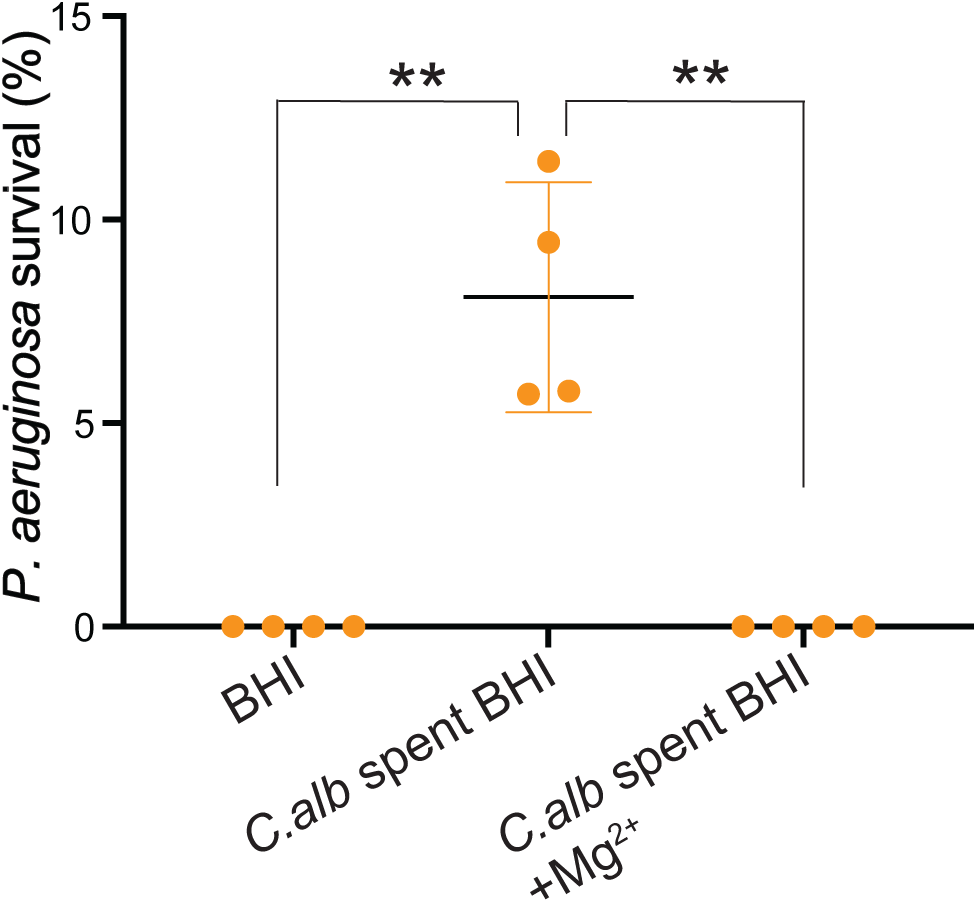
*C. albicans*-mediated Mg^2+^ sequestration confers resistance to polymyxin B by *P. aeruginosa*. The WT *P. aeruginosa* strain was highly susceptible to 2.5 μg/ml polymyxin B in BHI but became resistant in *C. albicans* spent BHI. Such resistance was restored to the WT level in fungal spent BHI supplemented with 10mM Mg^2+^. Mean ± std of bacterial survival from four biological replicates is shown (** *p*< 0.01, unpaired two-tailed Student’s *t*-test used).

**Figure S11.**
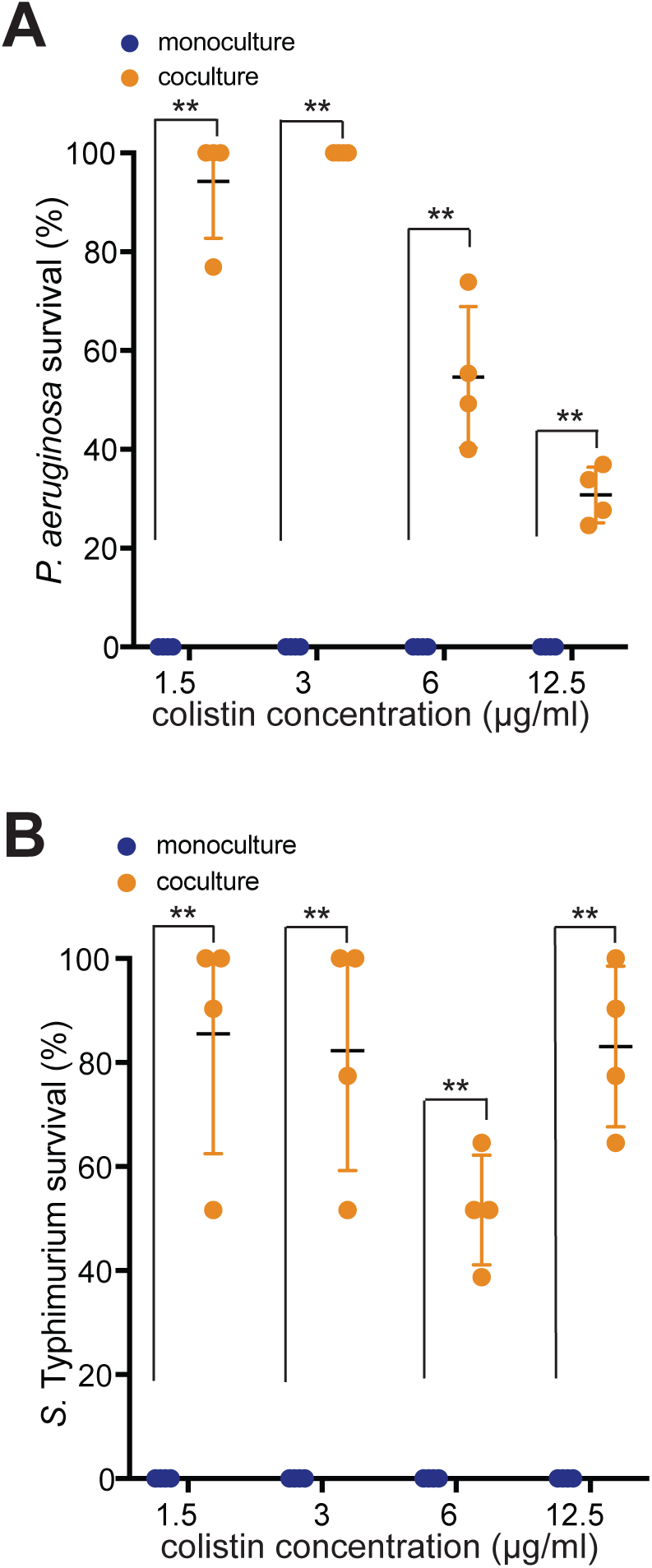
*C. albicans* protects both *P. aeruginosa* and *S.* Typhimurium from colistin at 37°C. (A) *P. aeruginosa* in co-culture with *C. albicans* was more resistant to colistin, as compared to monoculture, at 37°C (** *p*< 0.01, unpaired two-tailed Student’s *t*-test used). (B) *S.* Typhimurium in co-culture with *C. albicans* was more resistant to colistin, as compared to monoculture, at 37°C (** *p*< 0.01, unpaired two-tailed Student’s *t*-test used).

**Figure S12.**
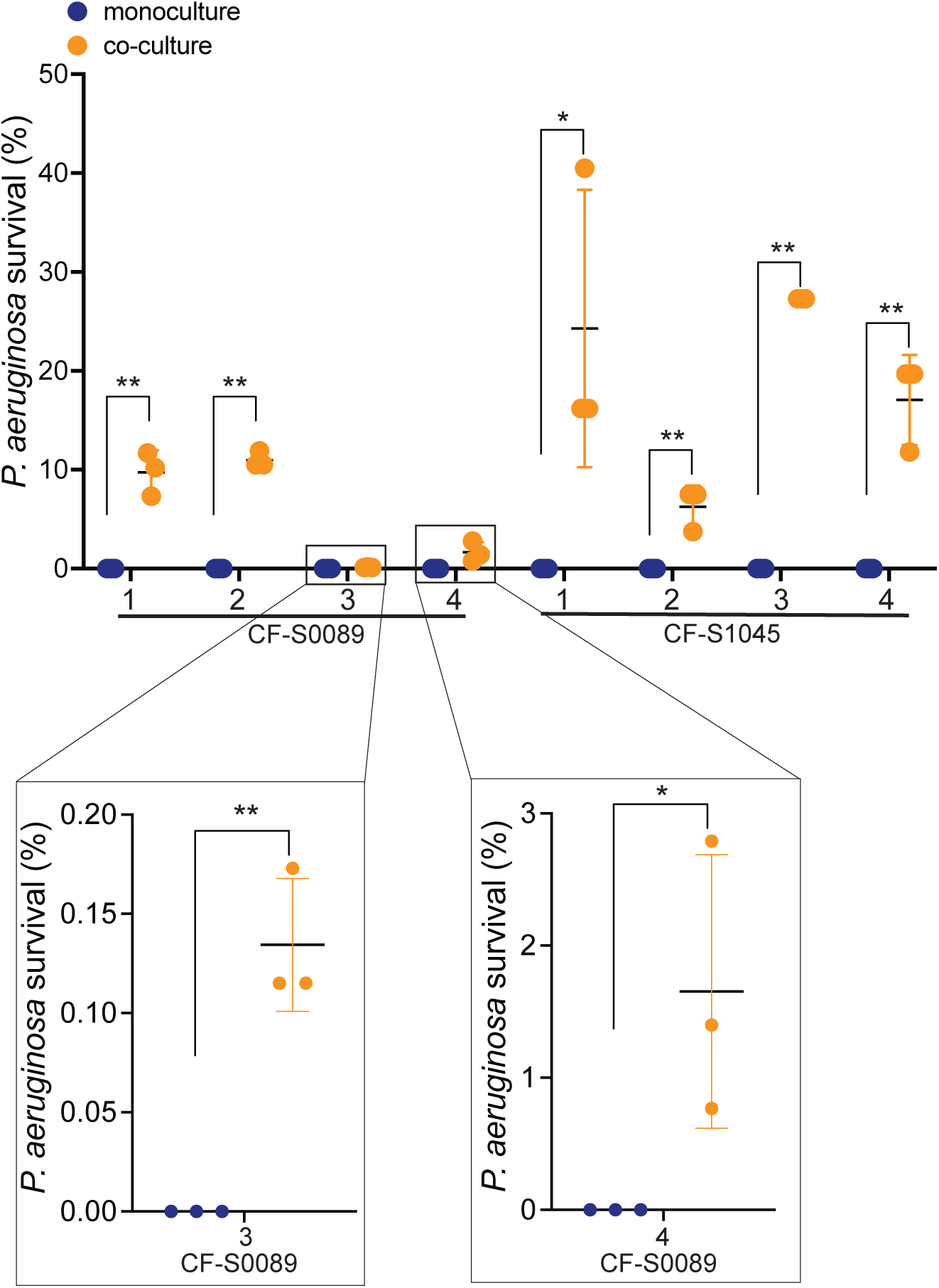
*C. albicans* protects *P. aeruginosa* CF isolates from colistin at 37^°^C. Eight isolates from two sputum samples (CF-S0089 and CF-S1045) were treated with 3 μg/ml colistin, either in monoculture or in co-culture with *C. albicans*, as indicated in colistin survival assay (See Methods). Bacterial survival is shown as the percentage of viable cells after the colistin treatment. Mean ± std of three biological replicates is shown (** *p* < 0.01, * *p* < 0.05, unpaired one-tailed Student’s *t* test used).

**Figure S13.**
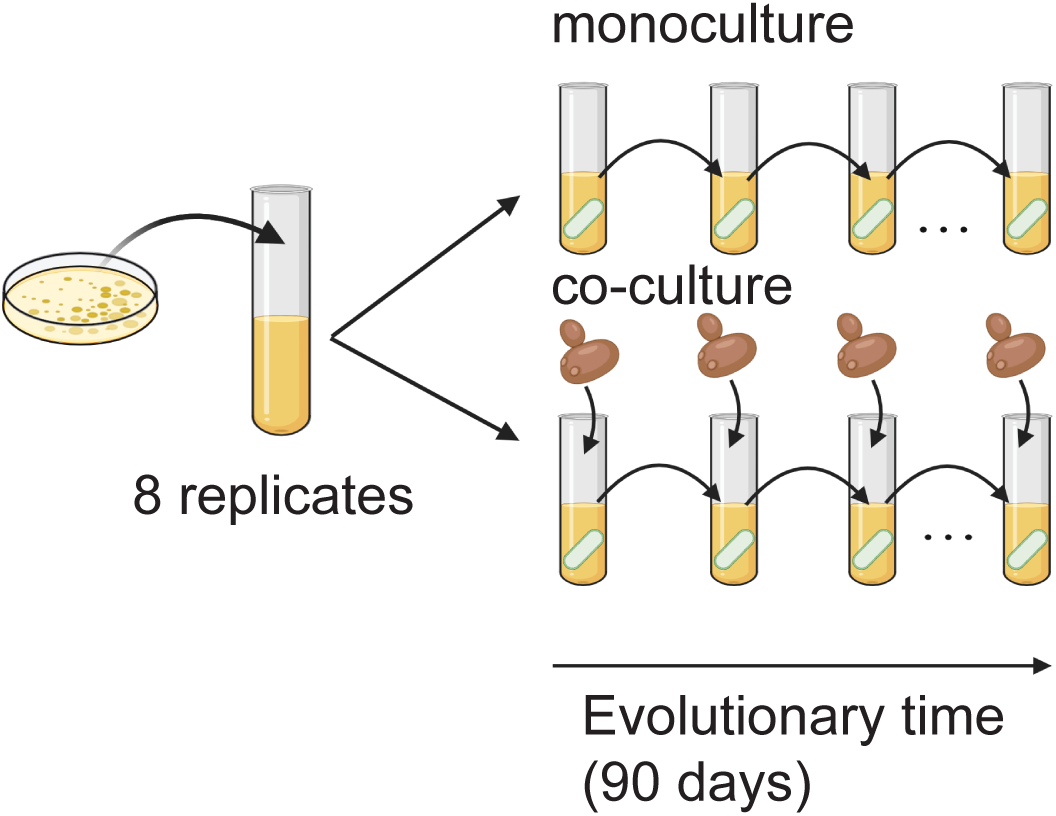
Evolution of *P. aeruginosa* in BHI. WT *P. aeruginosa* populations were passaged daily without colistin, to assess bacterial adaptation to BHI or co-culture with *C. albicans* in the absence of colistin. The diagram was generated using BioRender.

**Figure S14.**
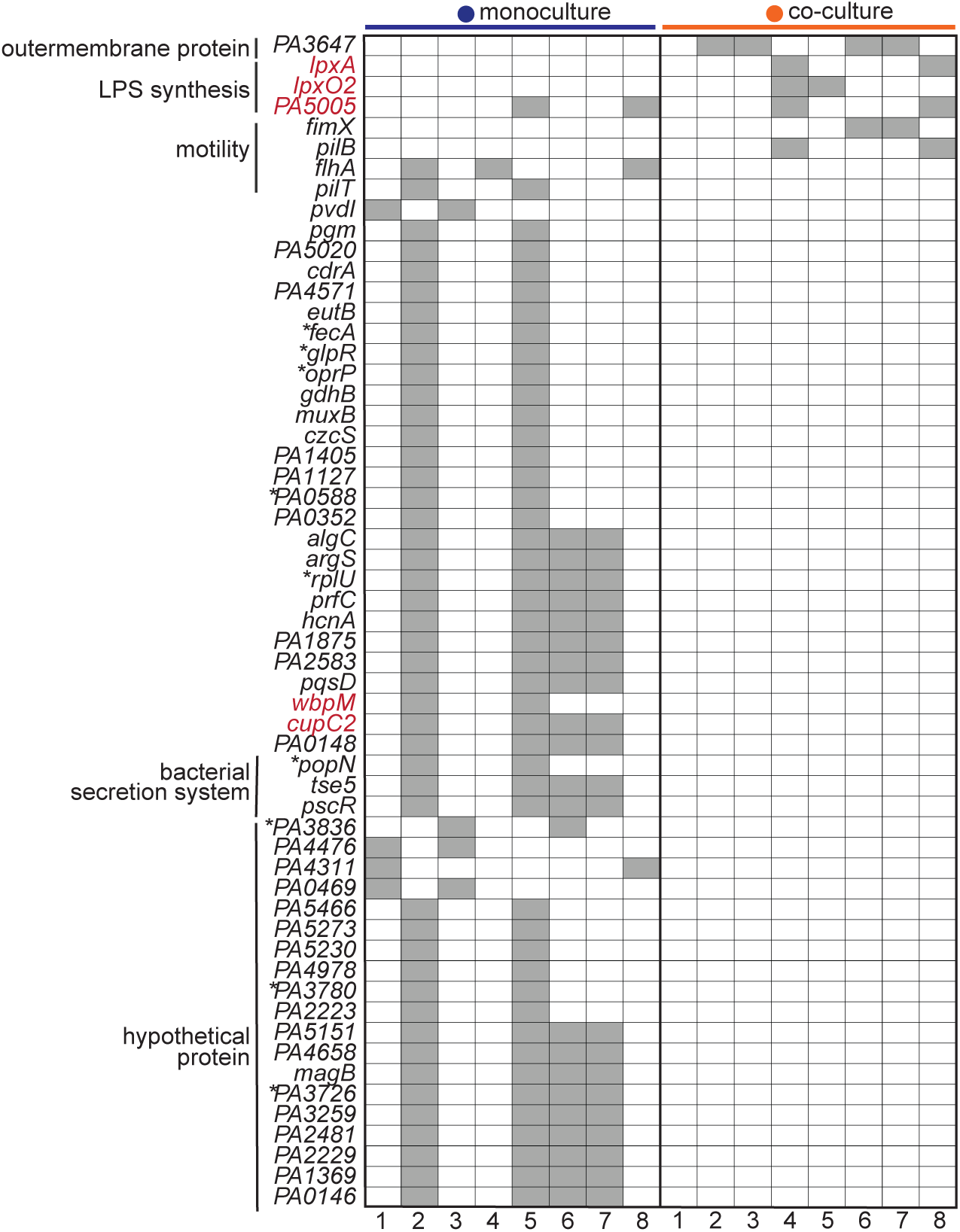
Other putative genetic targets for colistin resistance. Individual genes that have fixed mutations in more than two of colistin-resistant monoculture or co-culture-evolved populations are listed. Columns refer to individual evolved populations, and rows refer to mutated genes grouped by cellular process. Genes known to be involved in colistin resistance are labeled in red. Gray or blank squares indicate presence or absence of mutation, respectively (* indicates a mutation in the promoter region).

## Supplementary Tables

**Table S1.** *P. aeruginosa* differentially expressed genes in co-culture versus monoculture, in BHI or BHI supplemented with 10mM Mg^2+\^

**Table S2.** *P. aeruginosa* dispensable gene candidates identified in Tn-seq

**Table S3.** Fixed mutations in each colistin-resistant evolved population

**Table S4.** Fixed mutations in each control population without colistin treatment

**Table S5.** Bacterial and fungal strains used in this study

**Table S6.** Primers and plasmids used in this study

